# NAD(P)HX repair deficiency causes central metabolic perturbations in yeast and human cells

**DOI:** 10.1101/302257

**Authors:** Julia Becker-Kettern, Nicole Paczia, Jean-François Conrotte, Chenchen Zhu, Oliver Fiehn, Paul P. Jung, Lars M. Steinmetz, Carole L. Linster

**Affiliations:** Luxembourg Centre for Systems Biomedicine, University of Luxembourg, L-4362 Esch-sur-Alzette, Luxembourg; European Molecular Biology Laboratory (EMBL), Genome Biology Unit, 69117 Heidelberg, Germany; NIH West Coast Metabolomics Center, University of California Davis, Davis, CA 95616, USA; Stanford Genome Technology Center, Stanford University, Palo Alto, CA 94304, USA; Department of Genetics, Stanford University School of Medicine, Stanford, CA 94305, USA

**Author notes:** For correspondence: Carole L. Linster, Luxembourg Centre for Systems Biomedicine, University of Luxembourg, 7 Avenue des Hauts Fourneaux, L-4362 Esch-sur-Alzette, Luxembourg.

**Keywords:** metabolite repair, NAD(P)HX dehydratase, NAD(P)HX epimerase, *Saccharomyces cerevisiae*, inborn errors of metabolism.

## Abstract

NADHX and NADPHX are hydrated and redox inactive forms of the NADH and NADPH cofactors, known to inhibit several dehydrogenases *in vitro*. A metabolite repair system that is conserved in all domains of life and that comprises the two enzymes NAD(P)HX dehydratase and NAD(P)HX epimerase, allows reconversion of both the *S*- and *R*-epimers of NADHX and NADPHX to the normal cofactors. An inherited deficiency in this system has recently been shown to cause severe neurometabolic disease in children. Although evidence for the presence of NAD(P)HX has been obtained in plant and human cells, little is known about the mechanism of formation of these derivatives *in vivo* and their potential effects on cell metabolism. Here, we show that NAD(P)HX dehydratase deficiency in yeast leads to an important, temperature-dependent NADHX accumulation in quiescent cells with a concomitant depletion of intracellular NAD^+^ and serine pools. We demonstrate that NADHX potently inhibits the first step of the serine synthesis pathway in yeast. Human cells deficient in the NAD(P)HX dehydratase also accumulated NADHX and showed decreased viability. In addition, those cells consumed more glucose and produced more lactate, potentially indicating impaired mitochondrial function. Our results provide first insights into how NADHX accumulation affects cellular functions and pave the way for a better understanding of the mechanism(s) underlying the rapid and severe neurodegeneration leading to early death in NADHX repair deficient children.

## INTRODUCTION

A derivative of the central cofactor NADH, that differs from the normal cofactor by the absence of the typical absorbance maximum at 340 nm, had already been described to be formed in the presence of glyceraldehyde-3-phosphate dehydrogenase (GAPDH) in the 1950s by Edwin Krebs and colleagues [1]. Nuclear magnetic resonance studies revealed that this derivative, called NADHX, resulted from hydration of NADH on the 5,6-double bond of the nicotinamide cycle, leading to the formation of *S*- and *R*-epimeric forms ((6*S*)- or (6*R*)-6-hydroxy-tetrahydro-NAD; designated (*S*)-NADHX and (*R*)- NADHX hereafter) depending on the orientation of the hydroxyl group at the C-6 atom (compare Figure S2) [2]. Acidic conditions and the presence of high phosphate concentrations also stimulate NADHX formation and enhance further conversion of the latter to a third derivative (cyclic NADHX) [3]. Analogously, the phosphorylated cofactor NADPH can undergo hydration and cyclisation, resulting in all three forms of NADPHX [4]. An enzymatic activity catalyzing an ATP-dependent reconversion of NADHX back to NADH had been described in yeast extracts briefly after the discovery of these non-canonical metabolites [5] and more recently the existence of an S-epimer specific NADHX dehydratase was proposed [6]. In 2011, the gene encoding this enzyme has been identified (designated *NAXD* in humans, **EC 4.2.1.93**) and an additional gene was shown to encode an NAD(P)HX epimerase (*NAXE* in humans, **EC 5.1.99.6**); together those genes encode an NAD(P)HX repair system that is conserved in the great majority of living species [7]. Subsequent subcellular localization studies revealed that these enzymes are also targeted to multiple subcellular compartments including the cytosol and the mitochondria for the mouse and *Arabidopsis* dehydratases and epimerases as well as the endoplasmic reticulum for the mouse dehydratase and the chloroplasts for the *Arabidopsis* dehydratase and epimerase [8–10]. The NAD(P)HX dehydratase and epimerase are two members of a growing list of enzymes that have been recognized in recent years to participate in a process called metabolite repair or metabolite proofreading and in which a panoply of protective enzymatic activities are required to prevent the accumulation of non-canonical, potentially toxic metabolites that are formed continuously via enzymatic side reactions or spontaneous chemical reactions [11–14]. The NAXE protein has been reported by other investigators to play a role in cholesterol efflux and angiogenesis [15–17]; it remains unclear how this role could be reconciled with the metabolite repair function [18].

Despite the very high degree of conservation of the NAD(P)HX repair system and its presumed role in preserving active forms of the central cofactors NAD and NADP and/or preventing accumulation of toxic derivatives thereof, no overt growth phenotypes have been identified so far in high-throughput screens with *Escherichia coli* or yeast strains deleted for the NAD(P)HX repair genes [19, 20]. Similarly, no growth phenotype was observed in *Arabidopsis* NAD(P)HX dehydratase knockout lines maintained under standard conditions [8, 10]. In yeast, the NAD(P)HX dehydratase gene has, however, been shown to be induced by heat-shock and DNA damage (stress response element-regulated gene; [21]) and NAD(P)HX dehydratase deficient *Bacillus subtilis* mutants displayed prolonged lag phases and reduced growth rates when cultivated under potassium limitation [22]. In apparent contrast to these absent or subtle phenotypes in experimental model organisms, recent case reports from young children indicate a critical role for NAD(P)HX repair in the central nervous system. Spiegel *et al.* [23] described five siblings of a consanguineous family with a homozygous mutation in the human NAD(P)HX epimerase gene (*NAXE*). All patients presented with a severe leukoencephalopathy following trivial febrile illnesses during their first year of life. Disease progression included general weakness and loss of acquired motor skills within days until the occurrence of death at the age of 5 at the latest. In a second case report, Kremer and colleagues described very similar disease progressions with onset of symptoms after febrile infections and rapid deterioration of the health status for six children from four families carrying biallelic mutations in *NAXE* [24]. They also found elevated NADHX levels in fibroblasts derived from the patients. To date, no disease-causing mutations have been reported yet for the NAD(P)HX dehydratase gene (*NAXD*).

Despite the recent evidence for the existence of NAD(P)HX in plants and human cells [10, 24] and for the clinical importance of the NAD(P)HX repair system [23, 24], very little is known about the intracellular reactions leading to NAD(P)HX formation as well as the effects of NAD(P)HX accumulation on metabolic pathways and other cellular processes. The aim of this study was to analyze the consequences of NAD(P)HX repair enzyme deficiency in yeast and human cells on cell growth and viability, gene expression, and metabolism and to identify factors influencing intracellular NADHX formation. We were able to show intracellular NADHX accumulation in both yeast and human cells, most importantly after deletion of the NAD(P)HX dehydratase gene. In a yeast NAD(P)HX dehydratase knockout strain, NADHX accumulation was accompanied by a decrease in intracellular NAD^+^ and serine levels of up to 60% compared to a wild-type control strain. We discovered that NADHX is interfering with serine metabolism in yeast by inhibiting the first step in the *de novo* serine biosynthesis pathway from 3-phosphoglycerate. While yeast NAD(P)HX dehydratase knockout cells did not show slowed growth under the conditions tested here, knocking out the *NAXD* gene in human cells led to a significant decrease of cell viability after prolonged cultivation times. Increased glucose consumption and lactate production in the human *NAXD* knockout cells also provided a preliminary indication for perturbed mitochondrial function in those cells.

## RESULTS

### NAD(P)HX dehydratase deficiency leads to NAD(P)HX accumulation and NAD^+^ depletion in yeast

Given that little is known about the formation and effects/roles of NADHX and NADPHX in living cells, our first aim was to study the consequences of NAD(P)HX dehydratase and/or epimerase deficiency on the intracellular levels of normal and damaged forms of the nicotinamide nucleotide cofactors in the simple eukaryotic model organism *Saccharomyces cerevisiae*. Yeast deletion mutants lacking the NAD(P)HX dehydratase gene (*YKL151C*), the NAD(P)HX epimerase gene (*YNL200C*), or both genes were cultivated in minimal defined medium supplemented with 2% D-glucose along with the wild-type strain (all strains were isogenic to BY4741). Metabolites were extracted according to our protocol adapted from Sporty *et al.* [25] at different stages of growth for subsequent analysis of NAD^+^, NADH, and NADHX derivatives by HPLC-UV. Based on retention time and UV absorption spectrum comparison to standard compounds, we could detect (*S*)-, (*R*)- and cyclic NADHX in all cell extracts, independently from the strain and for most of the sampling points, even though for some samples in only very small amounts (Figure S1). Strikingly, in postdiauxic phase cells, (*S*)-, (*R*)-, and cyclic NADHX were found to accumulate to high levels in the *ykl151c*Δ (Figure S2) and *ykl151c*Δ*ynl200c*Δ strains (*ykl151c*Δ mutant vs. WT ratios > 40 for all NADHX derivatives), but not significantly (p > 0.05; apart from cyclic NADHX, p = 0.031) in the *ynl200c*Δ strain (*ykl151c*Δ mutant vs. WT ratios of 4.8, 1.4, and 2.8 for (*S*)-NADHX, (*R*)-NADHX, and cyclic NADHX, respectively) (Fig. 1A). The absence of significant NADHX accumulation in the *ynl200c*Δ strain (as also observed during exponential growth; not shown) suggested that, under the conditions tested, spontaneous intracellular epimerization between (*R*)- and (*S*)-NADHX and the presence of the NAD(P)HX dehydratase were sufficient to prevent NADHX from building up. Similar results were obtained for all four strains using an LC-MS method for NADH(X) analysis based on retention time and accurate mass for compound identification [10] (Figure S3A). Using this LC-MS method, NADPHX could also be detected (although with lower signal intensities and reproducibility) and was also shown to accumulate in the NAD(P)HX dehydratase deficient strains, but to a much lesser extent in the *ynl200c*Δ strain (Figure S3B). In contrast to the results obtained (for NADHX) by HPLC-UV, the cyclic NAD(P)HX derivatives were the predominant forms detected using the LC-MS method, which may at least partially be explained by slow cyclisation of (*S*)- and (*R*)-NAD(P)HX during the longer sample storage durations before LC-MS analysis, as also suggested previously [10]. Taken together, these results indicated that hydration of both NADH and NADPH can occur intracellularly and that the Ykl151c dehydratase is the main enzyme responsible to limit NAD(P)HX accumulation under standard cultivation conditions in yeast. Also, based on these results, we decided to focus for the rest of this study on the *ykl151c*Δ strain.

**Figure 1:**
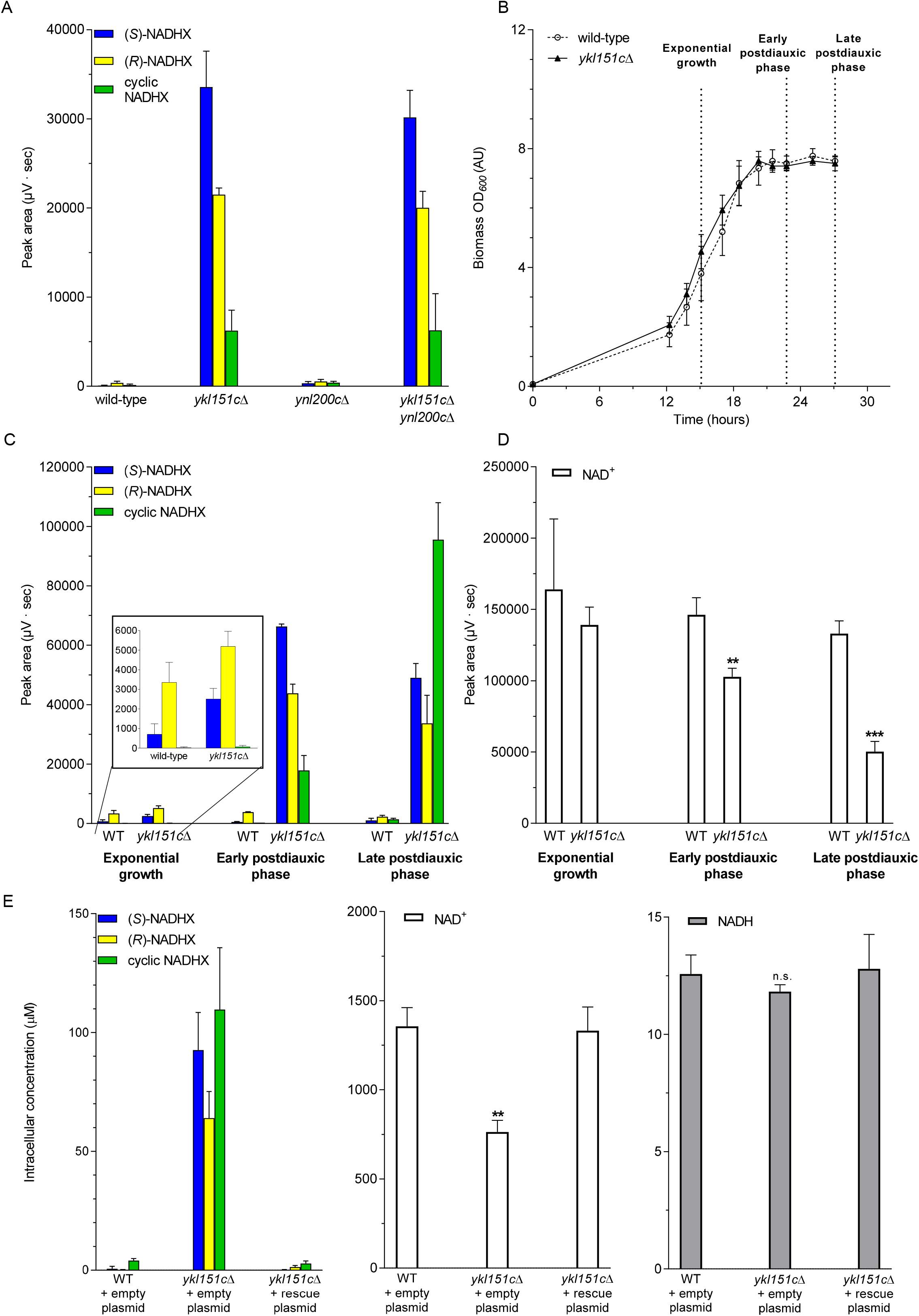
NADHX accumulates during the postdiauxic phase in yeast strains lacking the NAD(P)HX dehydratase Ykl151c. Yeast strains deleted for the *YKL151C* and/or *YNL200C* genes encoding NAD(P)HX dehydratase and NAD(P)HX epimerase, respectively, were cultivated in minimal defined medium containing uracil (A-D) or no uracil (E). NAD and NADHX derivatives were extracted at different stages of growth using methods adapted from Sporty *et al.* [25] (A, samples were extracted 1 hour after entering the postdiauxic phase; C, and D) or from Becker-Kettern *et al.* [38] (E, samples were extracted 5.5 hours after entering the postdiauxic phase) and measured by HPLC-UV (A, C, D, E). For panels A, C, and D, the biovolume was not determined and therefore peak areas are shown, whereas for panel E, the total biovolume of yeast cells was determined as described in the Methods section, allowing for calculation of intracellular concentrations. Growth was followed by measuring absorbance at 600 nm (B) and dashed lines indicate the sampling points for panels (C) and (D). All values shown are means and standard deviations of at least three independent replicates. All concentrations of (*S*)-, (*R*)-, and cyclic NADHX were significantly (p < 0.01) increased in postdiauxic phase in strains deleted for the *YKL151C* gene. During exponential growth only the S-NADHX concentration was significantly increased in the *ykl151c*Δ vs. wild-type strain (p = 0.014). In the *ynl200c*Δ strain, the only significantly changed NADHX derivative was the cyclic form (p = 0.031). For clarity, statistical significance is not indicated in the figure for the NADHX derivatives, but for NAD^+^ statistically significant differences compared to the wild-type strain are indicated by asterisks (*p < 0.05; **p < 0.01, ***p < 0.001; n.s., not significant).

As can be seen in Fig. 1B, the wild-type and *ykl151c*Δ strains presented identical growth properties under the cultivation conditions used (300 ml cultivations grown in minimal defined medium with 2% glucose in 2 liter flasks at 30 °C under continuous shaking). Similarly, no difference in growth rate or biomass yield between the two strains was found in more than 170 additional environmental conditions tested (in 96- and 384-well plate microcultivation format, shaking flasks, and/or agar plates) including the presence of various chemical stressors (e.g. methanol, ethanol, dithiothreitol, glycolaldehyde), exposure to different temperatures (continuously, at 25 to 37 °C; short-term, < 180 minutes, at 30 to 70 °C), and replacement of glucose with different carbon sources (e.g. glycerol, pyruvate, galactose; also in combination with other stresses). The list of all the environmental conditions tested is given in Table S1. Inoculation of main cultures from postdiauxic and stationary phase cultures instead of exponentially growing cultures also did not lead to a detectable growth phenotype of the *ykl151c*Δ strain compared to the wild-type strain (Table S1).

Based on our initial NAD(P)(H)(X) profiling results, showing, in contrast to the growth behavior, clear differences between the wild-type and *ykl151c*Δ strains, we next repeated those measurements in a time-course experiment where metabolites were extracted during exponential growth, at the entrance of postdiauxic phase, and 5.5 hours into postdiauxic phase (as shown in Fig. 1B). In the exponential growth phase, only very low levels of total NADHX (sum of (*S*)-, (*R*), and cyclic NADHX) were detected in both strains, but a slight increase was observed in the *ykl151c*Δ strain (3.5-fold for (*S*)-, 1.5-fold for (*R*)-, and 2.8-fold for cyclic NADHX, Fig. 1C). Upon entrance into postdiauxic phase, accompanied by glucose depletion and decreased cell division rate, similarly low NADHX levels were still found in the wild-type strain; in contrast, the total NADHX levels measured in the *ykl151c*Δ strain had increased more than 16-fold compared to exponential growth. In this same strain, it could also clearly be seen that the relative abundance of the different NADHX derivatives was growth phase dependent. Whereas the cyclic form of NADHX, which is derived from the *S*- and *R*-epimers of NADHX [7], represented a minor fraction of the total NADHX measured at the beginning of the postdiauxic phase, this derivative represented the predominant fraction later in postdiauxic phase (Fig. 1C and Fig. S2). While we currently ignore the precise reasons underlying this growth phase-dependent NADHX profile in the *ykl151c*Δ strain, decreased cell proliferation and switch from fermentative to respiratory metabolism may both contribute to NADHX accumulation becoming apparent only in postdiauxic phase.

The increase in NADHX levels was accompanied by a decrease in NAD^+^ levels in the *ykl151c*Δ strain (Fig. 1D and Fig. S2). While this decrease was only moderate and not statistically significant (p = 0.45) during exponential growth, a 30% and more than 60% decrease in NAD^+^ levels in the *ykl151c*Δ strain compared to the wild-type strain was measured in early and late postdiauxic phase, respectively.

The metabolic phenotypes (NADHX accumulation as well as NAD^+^ depletion) described above for the *ykl151c*Δ strain could be reverted by restoring *YKL151C* expression in this deletion mutant (Fig. 1E). While a *ykl151c*Δ strain transformed with an empty control vector accumulated 93 µM, 64 µM, and 110 µM of (*S*)-, (*R*)- and cyclic NADHX, respectively, the total NADHX levels measured in the wild-type control and rescue strains remained below 5 µM (Fig. 1E). Similarly, the more than 40% drop in NAD^+^ levels measured in the *ykl151c*Δ strain compared to the wild-type strain (from about 1.4 mM to less than 0.8 mM) was completely prevented in the rescue strain (Fig. 1E). These results confirmed that both NADHX accumulation and NAD^+^ depletion were caused specifically by the *YKL151C* gene deletion and not by a secondary mutation in another gene of the *ykl151c*Δ strain. No significant difference between the *ykl151c*Δ and wild-type strains was found for NADH levels (Fig. 1E and Table 1). NADHX repair deficiency thus seems to cause depletion of the NAD^+^ pool in the yeast cell, while NADH levels can be maintained by a presently unknown mechanism. This led to a decrease in the NAD^+^/NADH ratio in the *ykl151c*Δ strain which was most pronounced in the late postdiauxic phase (Table 1).

**Table 1:** NAD(P)HX dehydratase deficiency decreases intracellular NAD^+^ levels and thereby changes the cellular redox state. Wild-type and *ykl151c*Δ mutant strains were transformed either with an empty control plasmid or with a rescue plasmid encoding NAD(P)HX dehydratase (Ykl151c). Cells were cultivated in minimal defined medium lacking uracil for plasmid selection. NAD^+^ and NADH were extracted at the indicated stages of growth using a method adapted from Becker-Kettern *et al.* [38] and measured by HPLC-UV. All values shown are means and standard deviations of three independent replicates.

|  | [NAD<sup>+</sup>] (μM) | [NADH] (μM) | [NAD<sup>+</sup>]/[NADH] |
| --- | --- | --- | --- |
| *Exponential growth* |  |  |  |
| Wild-type + empty plasmid | 1280.4 ± 140.4 | 299.1 ± 33.8 | 4.3 ± 0.7 |
| Wild-type + rescue plasmid | 1257.6 ± 88.7 | 343.5 ± 24.4 | 3.7 ± 0.4 |
| *ykl151cΔ*+ empty plasmid | 1415.9 ± 141.2 | 397.8 ± 32.5 | 3.6 ± 0.5 |
| *ykl151cΔ*+ rescue plasmid | 1161.7 ± 74.9 | 346.1 ± 10.2 | 3.4 ± 0.2 |
| *Early postdiauxic phase* |  |  |  |
| Wild-type + empty plasmid | 1322.7 ± 29.0 | 79.9 ± 9.8 | 16.6 ± 2.1 |
| Wild-type + rescue plasmid | 1239.7 ± 67.5 | 143.9 ± 26.1 | 8.6 ± 1.6 |
| *ykl151cΔ*+ empty plasmid | 1052.7 ± 94.1 | 72.8 ± 8.0 | 14.5 ± 2.0 |
| *ykl151cΔ*+ rescue plasmid | 1217.5 ± 54.1 | 125.5 ± 5.6 | 9.7 ± 0.6 |
| *Late postdiauxic phase* |  |  |  |
| Wild-type + empty plasmid | 1356.3 ± 104.4 | 12.6 ± 0.8 | 107.9 ± 10.8 |
| Wild-type + rescue plasmid | 1462.7 ± 105.7 | 13.5 ± 1.3 | 108.0 ± 12.9 |
| *ykl151cΔ*+ empty plasmid | 762.9 ± 65.6 | 11.8 ± 0.3 | 64.5 ± 5.8 |
| *ykl151cΔ*+ rescue plasmid | 1331.5 ± 133.0 | 12.8 ± 1.5 | 104.1 ± 15.8 |

**Table 2:**
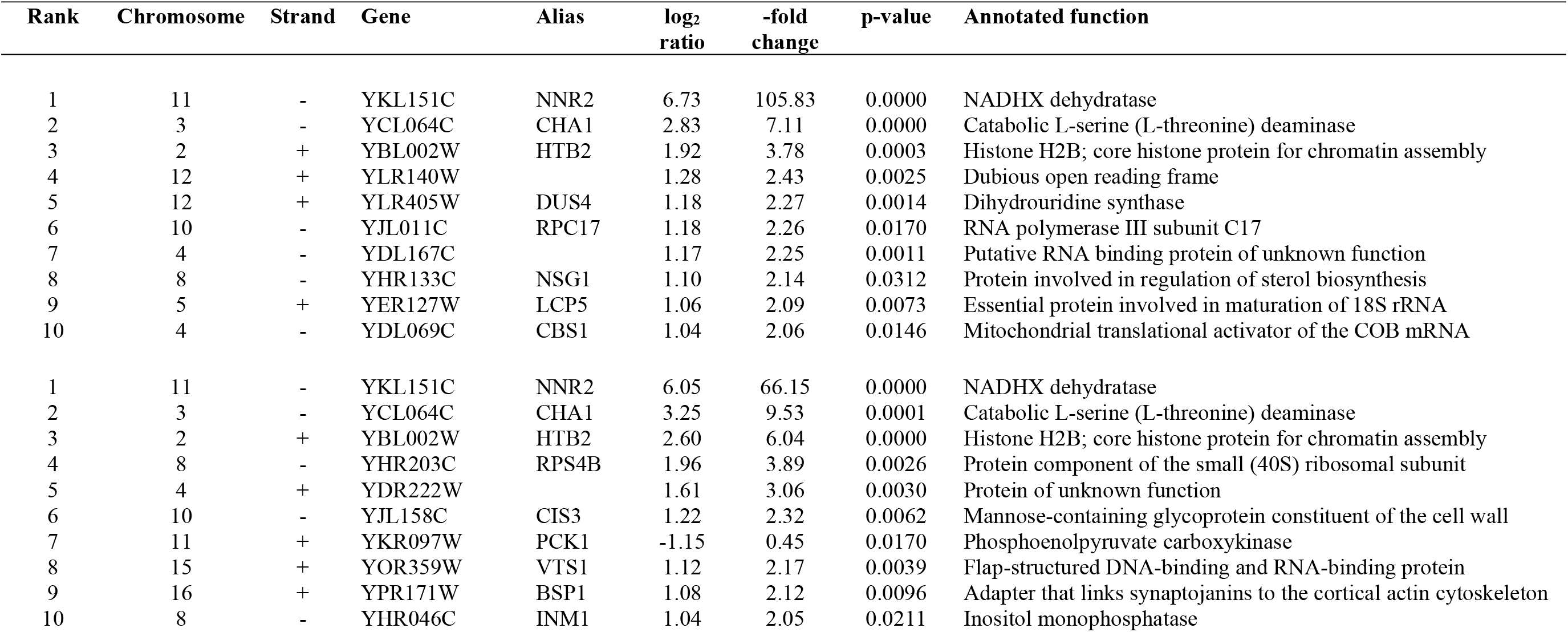
Top 10 significantly changed genes with most different expression levels between wild-type and ykl151cΔ strains. Genome-wide expression levels were determined in wild-type and *ykl151c*Δ strains using tiling array technology. Significantly changed genes (p < 0.05) were ranked according to absolute log2 ratios of wild-type over knockout, reflecting –fold changes between the two strains. The upper and lower halves of the table represent results obtained for the first sampling point in early postdiauxic phase and the second sampling point in late postdiauxic phase, respectively. The p-values were calculated based on background-corrected log2-expression values of three biological replicates for each strain using a Student’s t-test. The annotated gene functions were taken from the *Saccharomyces* Genome Database (https://www.yeastgenome.org/).

| Rank | Chromosome | Strand | Gene | Alias | $\log_2$<br>ratio | -fold<br>change | p-value | Annotated function |
| --- | --- | --- | --- | --- | --- | --- | --- | --- |
| 1 | 11 | - | YKL151C | NNR2 | 6.73 | 105.83 | 0.0000 | NADHX dehydratase |
| 2 | 3 | - | YCL064C | CHA1 | 2.83 | 7.11 | 0.0000 | Catabolic L-serine (L-threonine) deaminase |
| 3 | 2 | + | YBL002W | HTB2 | 1.92 | 3.78 | 0.0003 | Histone H2B; core histone protein for chromatin assembly |
| 4 | 12 | + | YLR140W |  | 1.28 | 2.43 | 0.0025 | Dubious open reading frame |
| 5 | 12 | + | YLR405W | DUS4 | 1.18 | 2.27 | 0.0014 | Dihydrouridine synthase |
| 6 | 10 | - | YJL011C | RPC17 | 1.18 | 2.26 | 0.0170 | RNA polymerase III subunit C17 |
| 7 | 4 | - | YDL167C |  | 1.17 | 2.25 | 0.0011 | Putative RNA binding protein of unknown function |
| 8 | 8 | - | YHR133C | NSG1 | 1.10 | 2.14 | 0.0312 | Protein involved in regulation of sterol biosynthesis |
| 9 | 5 | + | YER127W | LCP5 | 1.06 | 2.09 | 0.0073 | Essential protein involved in maturation of 18S rRNA |
| 10 | 4 | - | YDL069C | CBS1 | 1.04 | 2.06 | 0.0146 | Mitochondrial translational activator of the COB mRNA |
| 1 | 11 | - | YKL151C | NNR2 | 6.05 | 66.15 | 0.0000 | NADHX dehydratase |
| 2 | 3 | - | YCL064C | CHA1 | 3.25 | 9.53 | 0.0001 | Catabolic L-serine (L-threonine) deaminase |
| 3 | 2 | + | YBL002W | HTB2 | 2.60 | 6.04 | 0.0000 | Histone H2B; core histone protein for chromatin assembly |
| 4 | 8 | - | YHR203C | RPS4B | 1.96 | 3.89 | 0.0026 | Protein component of the small (40S) ribosomal subunit |
| 5 | 4 | + | YDR222W |  | 1.61 | 3.06 | 0.0030 | Protein of unknown function |
| 6 | 10 | - | YJL158C | CIS3 | 1.22 | 2.32 | 0.0062 | Mannose-containing glycoprotein constituent of the cell wall |
| 7 | 11 | + | YKR097W | PCK1 | -1.15 | 0.45 | 0.0170 | Phosphoenolpyruvate carboxykinase |
| 8 | 15 | + | YOR359W | VTS1 | 1.12 | 2.17 | 0.0039 | Flap-structured DNA-binding and RNA-binding protein |
| 9 | 16 | + | YPR171W | BSP1 | 1.08 | 2.12 | 0.0096 | Adapter that links synaptojanins to the cortical actin cytoskeleton |
| 10 | 8 | - | YHR046C | INM1 | 1.04 | 2.05 | 0.0211 | Inositol monophosphatase |

### Effect of temperature on intracellular NADHX levels in yeast

NAD(P)HX formation from NAD(P)H is known to be accelerated by increased temperature and by the presence of glyceraldehyde-3-phosphate dehydrogenase (GAPDH, **EC 1.2.1.12**) *in vitro* [1, 2, 7]. In addition, Ykl151c has been reported as a stress induced protein [21] and the expression of both the *YKL151C* and *YNL200C* genes is increased upon exposure to elevated temperatures [26]. We therefore wanted to test whether changes in temperature and expression of GAPDH also modulate NADHX levels in the living cell.

As shown in Fig. 2A, shifting the cultivation temperature from 30 °C to 37 °C led to an about 1.5-fold increase for both the (*S*)- and (*R*)-NADHX levels in *ykl151c*Δ cells. No significant decrease in NADHX levels was observed when cells were exposed to a lower temperature (25 °C). Here, it should be noted that in wild-type extracts prepared shortly after the diauxic shift, a compound co-eluting with (*R*)-NADHX (but with a different UV absorption spectrum) interfered with our (*R*)-NADHX measurements. In the *ykl151c*Δ strain, however, the predominant compound eluting at this retention time was (*R*)-NADHX, as confirmed by the UV absorption spectrum. Assuming that the contaminant was present in similar amounts in both strains, (*S*)- and (*R*)-NADHX were detected in the *ykl151c*Δ mutant strain in a ratio (65:35) consistent with previously reported *in vitro* results [2], when considering corrected (*R*)-NADHX levels. The increase in NADHX accumulation at 37 °C in the *ykl151c*Δ strain was accompanied by a more pronounced decrease in intracellular NAD^+^ levels in the *ykl151c*Δ mutant strain compared to the wild-type strain at this higher temperature (Fig. 2B), which further supports that repair of damaged NADH is important for maintaining the functional pool of NAD^+^. The NADHX repair capacity of the yeast cells did not seem to be exceeded under the conditions tested, as no increase of the NADHX levels was observed in the wild-type cells even at 37 °C, possibly also due to the induction of the NADHX repair enzymes by the heat stress. Our results in the yeast model corroborate recent observations in patients deficient in the *NAXE* gene (encoding the NAD(P)HX epimerase) [23, 24], where severe disease onset followed in general trivial febrile episodes, indicating that temperature plays a pivotal role in modulating intracellular NADHX levels especially when the NADHX repair system is deficient.

**Figure 2:**
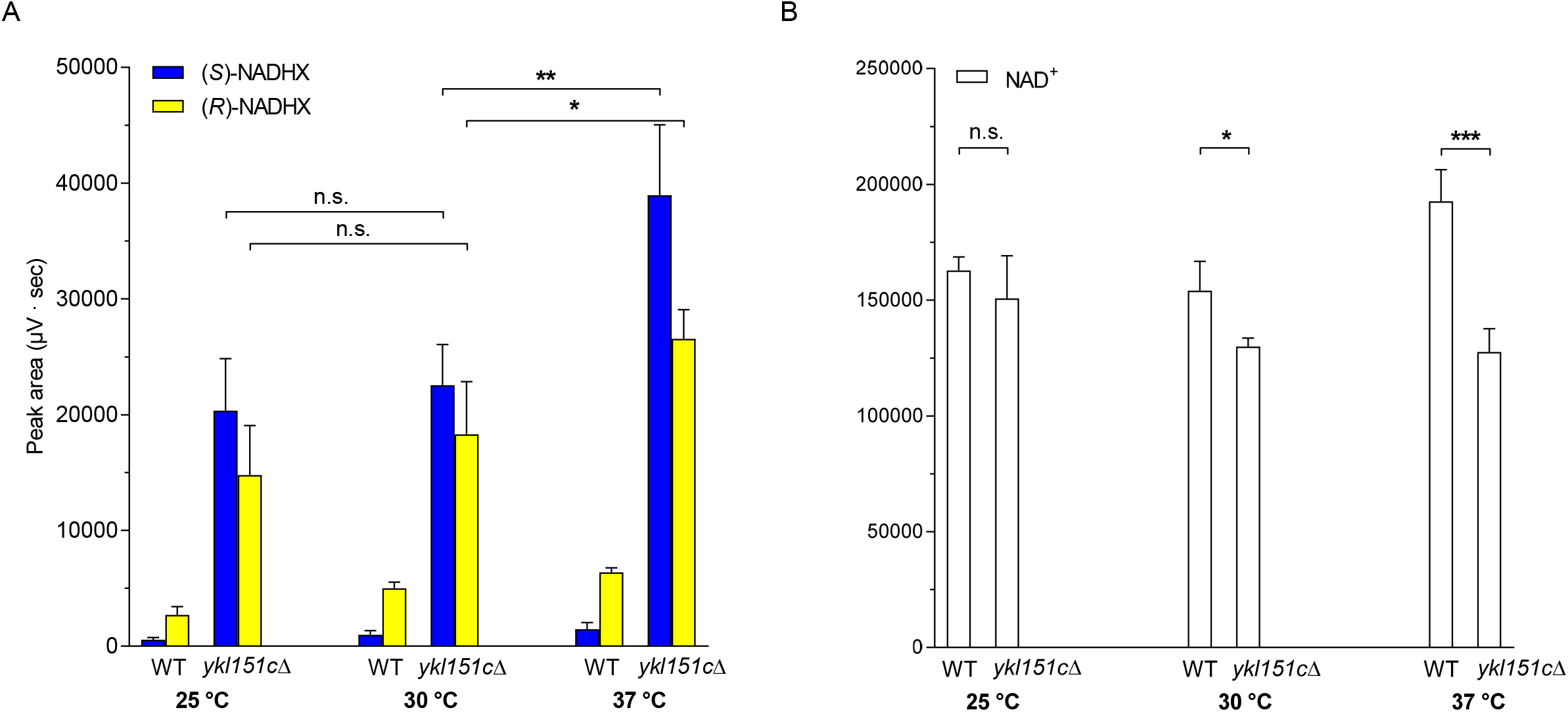
NADHX levels increase at elevated temperature. (A, B) Wild-type and *ykl151c*Δ mutant strains were cultivated in minimal defined medium with 2% glucose at 25 °C, 30 °C, or 37 °C. Metabolites were extracted around 2 hours into postdiauxic phase using a method adapted from [25]. NAD^+^ and NADHX derivatives were analysed by HPLC-UV. The values shown are means and standard deviations of at least three biological replicates. Statistical significance was calculated to compare NADHX levels in the *ykl151c*Δ mutant at different temperatures (A) or NAD levels between wild-type and *ykl151c*Δ mutant using an equal variance, unpaired Student’s t-test (B).

To test whether GAPDH serves as an *in vivo* source for NADHX, as suggested by previous *in vitro* observations [1], we tried to individually overexpress the three yeast GAPDH isoforms (encoded by the *TDH1*, *TDH2*, and *TDH3* genes in yeast), using a plasmid-based system, in the *ykl151c*Δ background. The Tdh isoforms display very similar amino acid sequences and comparable kinetic properties [27]. However, in exponentially growing cells, Tdh2 and Tdh3 are the major isoforms, accounting for almost 90% of the total GAPDH activity [27], whereas Tdh1 represents the predominant isoform in stationary phase [28]. Tdh1, but not Tdh2 or Tdh3, was also reported to be induced upon accumulation of cytosolic NADH [29]. For reasons that remain unclear, we were unable, however, despite testing the three yeast TDH isoforms, two different overexpression systems (based either on the pAG416-GPD-ccdB vector (Addgene ID #14148, constitutive GPD promoter, low copy number, *URA3* for plasmid selection) or the pAG426-GAL1-ccdB vector (Addgene ID #14155, inducible GAL1 promoter, high copy number, *URA3* for plasmid selection), and different cultivation conditions (minimal medium and synthetic complete medium; presence of 2% galactose in case of the GAL1 vector), to induce a robust increase in expression of this enzyme in our yeast strain (as tested by Western blotting) and did therefore not further pursue our attempts to validate GAPDH as a potentially major source of intracellular NADHX production. The relative contribution, if any, of Tdh isoforms to endogenous NADH hydration could also be explored using the opposite approach, i. e. deletion of one or more of the *TDH* genes in the *ykl151c*Δ background. Other NAD-dependent enzymes should, however, be considered as well as potential endogenous sources for NADHX in future work.

### Impact of intracellular NADHX accumulation on gene expression and amino acid levels in yeast

Given the central roles of the NAD(P) cofactors in metabolism, but also the regulation of gene expression via NAD-dependent histone deacetylases (sirtuins), the above described metabolic changes caused by NAD(P)HX repair deficiency could be expected to affect a panoply of downstream cellular processes. More specifically, NADHX and NADPHX have already been shown to potently inhibit, *in vitro*, glycerol-3-phosphate dehydrogenase (**EC 1.1.1.8**) [31] and the pentose phosphate pathway enzymes glucose-6-phosphate dehydrogenase (**EC 1.1.1.49**) and 6-phosphogluconate dehydrogenase (**EC 1.1.1.44**) [4], respectively. Such metabolic blockages could, in addition to changes in metabolite pool levels and pathway fluxes trigger (compensatory) changes in (metabolic) gene expression in the living cell.

In order to explore more globally if and how NAD(P)HX repair deficiency impacts gene expression, we performed high-density oligonucleotide tiling array analyses [32–34] to compare the transcriptome of the *ykl151c*Δ mutant to the corresponding wild-type strain, at two time points of postdiauxic phase where we had previously demonstrated NADHX accumulation and NAD^+^ depletion (see Figs. 1B, 1C, and 1D). 271 and 467 genes (Stable Unannotated Transcripts and Cryptic Unstable Transcripts [33] were not taken into account for the presented data analysis) showed significantly changed expression levels (p < 0.05 based on an equal variance, unpaired Student’s t-test) in the *ykl151c*Δ strain compared to the wild-type strain at the first and second time point, respectively (Fig. 3A and Table S2). Gene ontology (GO) enrichment analyses, performed separately for the significantly up- and downregulated genes at both time points using the GOTermFinder tool [35], failed to identify a specific biological process, molecular function or cellular component significantly linked to NAD(P)HX repair deficiency. We also carried out a differential pathway analysis on metabolic pathways from the YeastCyc database (https://yeast.biocyc.org) for both time points, adjusting the significance scores for multiple hypothesis testing using the Benjamini-Hochberg method. Including only pathways with 10 or more mapped genes, the ‘lipid-linked oligosaccharide biosynthesis’ pathway (adjusted p-value = 0.02) and the pentose phosphate pathway (adjusted p-value = 0.06, i.e. just above the statistical significance threshold) were found to be downregulated in the *ykl151c*Δ mutant strain. Ranking the significantly (p < 0.05) changed genes according to the absolute log2 ratio of wild-type over *ykl151c*Δ mutant, the three most differentially expressed genes (all downregulated in the *ykl151c*Δ strain) at both time points were *YKL151C* (i.e. the gene deleted in the *ykl151c*Δ strain), *CHA1,* and *HTB2* (Table 1). Given the high wild-type/*ykl151c*Δ –fold changes, at both time points tested, for *CHA1* (> 7-fold), encoding a serine (threonine) deaminase (**EC 4.3.1.17**), and for *HTB2* (> 3.8-fold), encoding the core histone protein H2B, we decided to focus on these genes for further investigations.

**Figure 3:**
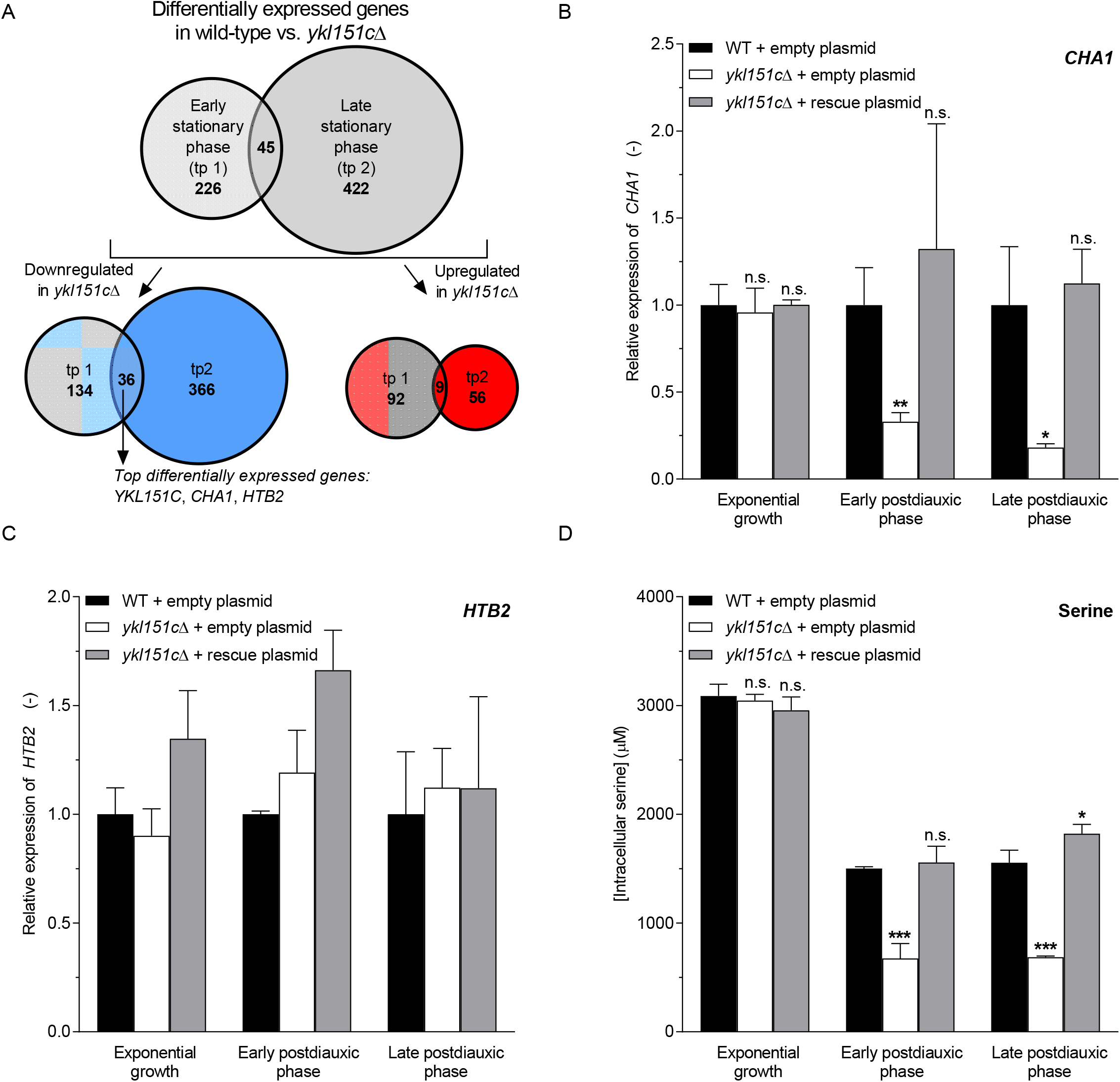
NAD(P)HX repair deficiency perturbs yeast serine metabolism. (A) Genes with significantly (p < 0.05) different expression levels between wild-type and *ykl151c*Δ mutant strains were identified at two sampling points in early (tp1) and late (tp 2) postdiauxic phase (grey circles, upper part) using transcriptomics analyses based on tiling arrays. Blue and red circles represent the number of genes that are downregulated and upregulated, respectively, in the *ykl151c*Δ versus wild-type strains. (B-D) Wild-type and *ykl151c*Δ mutant strains were transformed with an empty control plasmid or with a rescue plasmid containing the *YKL151C* coding sequence and cultivated in minimal defined medium with 2% glucose and without uracil. Transcript levels were determined at the indicated growth stages by qRT-PCR for the *CHA1* (B) and *HTB2* (C) genes. Fold changes were calculated using the 2^−ΔΔ*CT*^ method and *ALG9* as the reference gene. (D) Intracellular serine concentrations were determined at the same growth stages using LC-MS/MS. All values shown in panels B—D are means and standard deviations of three biological replicates. Statistically significant differences compared to wild-type control were determined as described in the Methods section (D) or applying an unpaired Student’s t-test and correcting for multiple comparisons using the Holm-Sidak method (B).

Decreased *CHA1* expression in the *ykl151c*Δ mutant was confirmed by using a qPCR approach (Fig. 3B). Interestingly, this downregulation was only observed during the two postdiauxic phase time points analyzed (3-fold and 5.5-fold decrease of *CHA1* expression at the early and late postdiauxic phase time point, respectively) and not during exponential growth (Fig. 3B), correlating therefore perfectly with the dynamics of NADHX accumulation observed previously in the *ykl151c*Δ mutant (see Fig. 1C). In the rescue strain, wild-type levels of *CHA1* were detected at all time points analyzed, confirming that the downregulation of this gene is specifically induced by the *YKL151C* gene deletion (Fig. 3B). By contrast, for reasons that remain uncertain, the *HTB2* downregulation found by the tiling array analyses in the *ykl151c*Δ mutant could so far not be validated by qPCR (Fig. 3C) and we therefore did not pursue our investigations on a possible link between the *HTB2* and *YKL151C* genes further in this study.

Because *CHA1*, a deaminase involved in serine and threonine catabolism [36], showed strongly decreased expression levels in the *ykl151c*Δ mutant, we wanted to investigate whether amino acid levels in general, and serine and threonine levels more specifically, were modified in this strain. Among the 20 amino acids measured, 5 showed significant (p < 0.05) differences in the *ykl151c*Δ strain versus wild-type strains (> 10% decrease) in early postdiauxic phase that were at least partially rescued upon re-expression of *YKL151C* in the *ykl151c*Δ background (Table S3). While the changes measured for glycine, asparagine, lysine, and proline were only moderate (< 20% decrease in *ykl151c*Δ versus wild-type strains), a more than 50% decrease in serine levels was measured in the *ykl151c*Δ mutant (Fig. 3D). A similar serine depletion was measured later in postdiauxic phase, while no difference in serine levels between wild-type and *ykl151c*Δ strains was detected in exponentially growing cells, mirroring again perfectly the growth stage dependent effects of *YKL151C* deletion on both NADHX levels and *CHA1* expression and suggesting a causal link between these molecular phenotypes. Wild-type serine levels were measured in the rescue strain at all time points (Fig. 3D) confirming that also serine depletion is specifically mediated by *YKL151C* deletion.

### NADHX potently inhibits the first step of the serine synthesis pathway in yeast

The main metabolic pathway for serine synthesis in yeast growing on glucose proceeds from the glycolytic intermediate 3-phosphoglycerate (3PGA) via oxidation to 3-phosphohydroxypyruvate (3PHP) by Ser3 or Ser33, followed by transamination to 3-phosphoserine (3PSer) by Ser1, and eventually dephosphorylation by Ser2 [37] (Fig. 4A, upper panel). Ser3 and Ser33 are two isozymes, currently annotated as NAD^+^-dependent 3PGA dehydrogenases (**EC 1.1.1.95**) based on sequence similarity with established 3PGA dehydrogenases from other species and serine auxotrophy of a *ser3*Δ*ser33*Δ double mutant [37]. As they catalyze the only supposedly NAD(P)-dependent step in this metabolic pathway, they appeared as the most plausible targets for an inhibitory effect of NADHX which could mediate the observed decrease in serine levels in the *ykl151c*Δ mutant. To test this hypothesis, we wanted to assay the enzymatic activity of recombinant Ser3 and Ser33 *in vitro* in the absence or presence of purified NADHX. As described in a previous publication, while we could detect an NADH-dependent α-ketoglutarate (αKG) reductase activity (producing D-2-hydroxyglutarate) with the purified yeast Ser3 and Ser33 enzymes, detecting the presumed physiological activity (NAD-dependent oxidation of 3PGA) proved much more difficult if not impossible [38].

**Figure 4:**
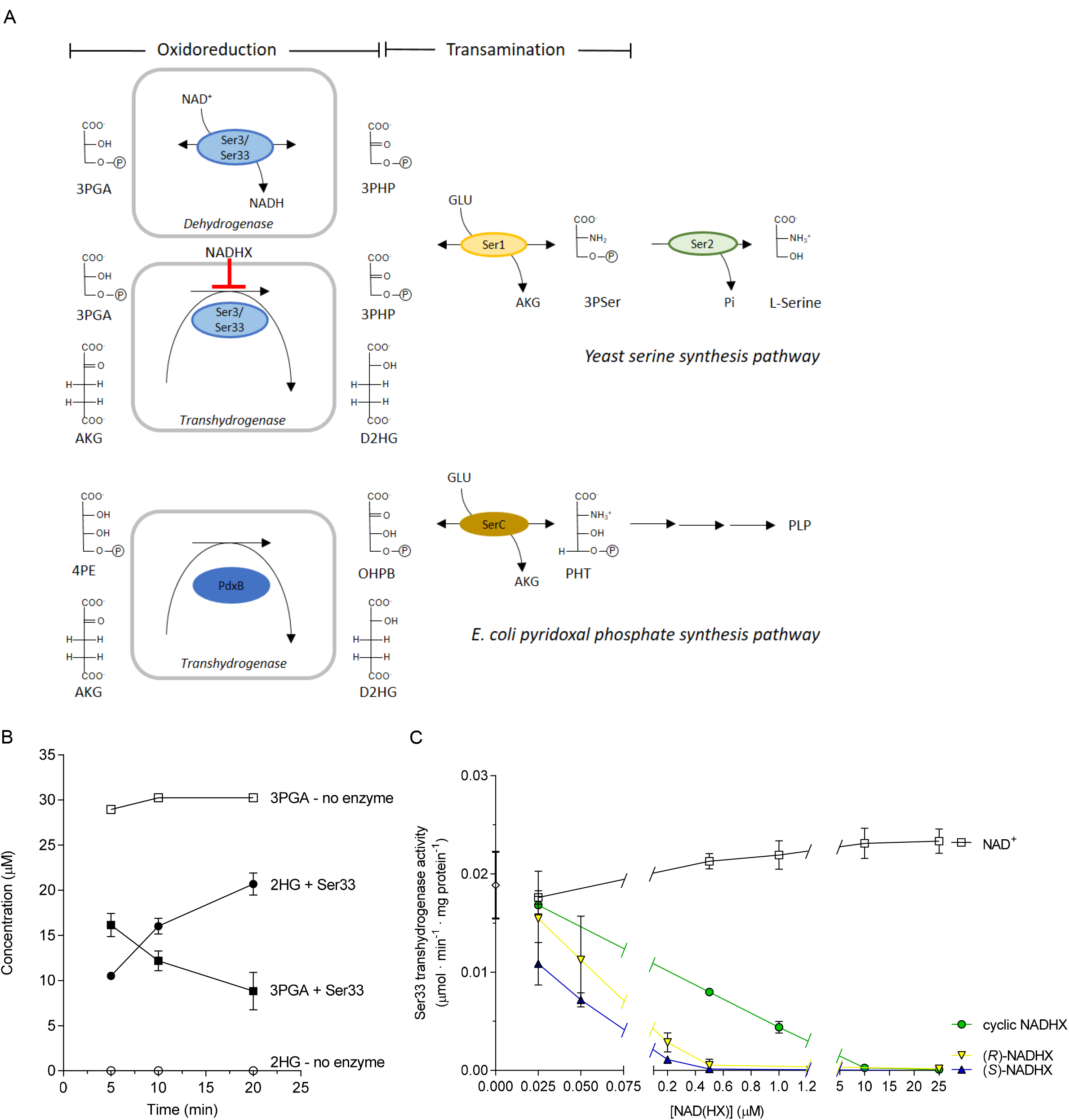
NADHX inhibits the first step of the main serine synthesis pathway in yeast. (A) Schematic overview of the serine synthesis pathway in yeast from the glycolytic intermediate 3PGA. Ser3 and its isoform Ser33, catalysing the first step of this pathway and currently annotated as 3PGA dehydrogenases (upper scheme), were found in this study to act as transhydrogenases (middle scheme), oxidizing 3PGA to 3PHP in the first half-reaction and reducing AKG to D2HG in the second half-reaction. The *E. coli* PLP synthesis pathway is represented (lower scheme) to highlight the striking parallels between the first two steps in this pathway and the serine synthesis pathway. (B) Purified recombinant Ser33 was incubated at 30 °C in a mixture containing 3PGA, AKG, purified recombinant human phosphoserine transaminase, glutamate, and PLP. The 3PGA consumption and concomitant 2HG formation measured by LC-MS/MS after incubation in a reaction mixture lacking externally added NAD^+^ demonstrate transhydrogenase activity. (C) Effect of NAD^+^, (*S*)-NADHX, (*R*)-NADHX and cyclic NADHX at the indicated concentrations on the transhydrogenase activity of Ser33. The values shown are means ± standard deviations (n = 3), except the no enzyme control incubation in (B), where single measurements are shown. AKG, α-ketoglutarate; D2HG, D-2-hydroxyglutarate; GLU, L-glutamate; 2HG, 2-hydroxyglutarate; OHPB, 2-oxo-3-hydroxy-4-phosphobutanoate; 4PE, 4-phospho-D-erythronate; 3PGA, 3-phosphoglycerate; 3PHP, 3-phosphohydroxypyruvate; PHT, 4-phosphohydroxy-L-threonine; PLP, pyridoxal-5’-phosphate; 3PSer, 3-phosphoserine.

This apparent lack of detectable 3PGA dehydrogenase activity of the Ser3 and Ser33 enzymes was reminiscent of the difficulties that other investigators had reported while trying to assay an NAD-dependent oxidation of 4-phosphoerythronate (structurally very close to 3PGA, with just one additional secondary alcohol group in the carbohydrate backbone) with the *Escherichia coli* PdxB enzyme (**EC 1.1.1.290**) [39]. PdxB catalyzes the second step of the pyridoxal phosphate synthesis pathway in *E. coli*, with the subsequent step consisting in a glutamate dependent transamination, just as the reaction following 3PGA oxidation in the serine synthesis pathway (Fig. 4A). It is also of note that *E. coli* PdxB shows 27% sequence identity with the yeast Ser3 and Ser33 proteins and that the latter are the best protein hits when blasting the *E. coli* PdxB sequence against the *S. cerevisiae* genome. It was eventually shown that PdxB is actually a nicotino-enzyme that tightly binds the NAD(H) cofactor and that depends on the presence of α-ketoacid acceptors (α-ketoglutarate, oxaloacetate, or pyruvate) to recycle enzyme-bound reduced NADH after 4-phosphoerythronate oxidation [39]. Taken together, these observations led us to test whether Ser3 and Ser33 act as transhydrogenases to oxidize 3PGA to 3PHP with a concomitant reduction of α-ketoglutarate to 2-hydroxyglutarate, rather than acting as dehydrogenases depending on exogenous NAD^+^ (Fig. 4A). Indeed, in a coupled end-point assay including the Ser3 or Ser33 enzyme as well as the human 3-phosphoserine aminotransferase homolog PSAT1 (**EC 2.6.1.52**; to pull the thermodynamically unfavorable 3PGA oxidation reaction [40]), we were able, using LC-MS/MS for compound identification and quantification, to demonstrate a stoichiometrically balanced consumption of 3PGA and formation of 2-hydroxyglutarate in the presence of α-ketoglutarate and in the absence of externally added NAD^+^ (Fig. 4B). A more detailed characterization of this newly identified enzyme activity and its significance for yeast central carbon metabolism will be published in a separate article as it goes beyond the scope of the present study.

Having found conditions under which we could assay 3PGA oxidation by Ser3 and Ser33 *in vitro*, we were now in a position to test whether NADHX can inhibit the first step of the serine synthesis pathway in yeast. Purified (*S*)-, (*R*)-or cyclic NADHX were added to the Ser3/Ser33 transhydrogenase assay at a final concentration of up to 25 µM. Specific activities were calculated based on 2-hydroxyglutarate formation measured by LC-MS/MS. As shown in Fig. 4C, the *S*- and *R*-epimers of NADHX and to a slightly lesser extent cyclic NADHX potently inhibited Ser33 activity. Complete inhibition of the latter was reached with 0.5 µM (*S*)- or (*R*)-NADHX, while 10 µM cyclic NADHX were required to fully block this activity. By contrast, a modest stimulation of up to 1.2-fold of the activity was observed in the presence of NAD^+^; this effect may be explained by a gradual partial release of the bound NAD cofactor from the purified enzyme during incubation at 30 °C. It should be noted that during these enzyme assays performed at neutral pH, a certain interconversion of the *S*- and *R*-epimers of NADHX is expected due to a spontaneous epimerization reaction [7]. Nevertheless, the curves obtained when adding purified (*S*)- or (*R*)-NADHX to the assays suggest that both epimers exert strong inhibitory effects, while cyclic NADHX is a less potent inhibitor. Similar results were found with the second isoenzyme Ser3 when testing the effects of (*S*)-, (*R*)-, or cyclic NADHX as well as NAD^+^ on the 3PGA-α-ketoglutarate transhydrogenase activity (data not shown).

Taken together our observations suggest that, in the *ykl151c*Δ mutant, the high intracellular NADHX levels measured in postdiauxic phase cells strongly inhibit the first step of the 3PGA serine synthesis pathway, leading to decreased intracellular levels of this amino acid. Reduced serine levels may in turn explain the lower *CHA1* transcript levels measured in the *ykl151c*Δ mutant, as expression of this gene is known to be regulated by serine availability [41, 42].

### Human cells are more sensitive to intracellular NADHX accumulation than yeast cells

The metabolic perturbations observed upon NAD(P)HX dehydratase deletion in yeast did not lead to any detectable growth phenotype under the cultivation conditions that we have tested so far. This is in apparent contrast with recent reports, where mutations in the other NAD(P)HX repair enzyme (NAD(P)HX epimerase encoded by the *NAXE* gene in humans) led to severe neurometabolic symptoms and early death in affected children [23, 24]. We therefore proceeded to investigate human cells deficient in the NAD(P)HX repair enzymes. We used HAP1 haploid cell lines that had been genetically modified by the CRISPR/Cas9 technology to disrupt the human homologs of the yeast *YKL151C* or *YNL200C* genes, i.e. *NAXD* or *NAXE*, respectively. Gene disruption was verified by sequencing of the corresponding genes using genomic DNA extracted from the cells. *In silico* translation from the residual gene sequences predicted very early truncation and therefore most likely nonfunctional expression products for both genes. In the *NAXD* KO cell line, the mutated *NAXD* gene encodes a truncated protein containing the first 126 amino acids (out of the 329 amino acids of the native mitochondrial isoform) followed by 9 non-native amino acids. In the *NAXE* KO cell line, expression of a truncated protein comprising the first 83 of the 288 amino acids of the native mitochondrial protein, followed by 4 non-native amino acids, can be predicted.

Wild-type and KO HAP1 cells were cultivated in 6-well plates for five days and cell number and viability were determined daily. After 72 hours of cultivation, a total number of about 3 million cells per well was reached for each cell line, with an average viability higher than 98% (Fig. 5A). Strikingly, while the cell number continued to steadily increase thereafter for the control and *NAXE* KO cell lines, reaching more than 12 million viable cells/well after 120 hours of cultivation, this increase was much more modest for the *NAXD* KO cell line (about 6.6 million viable cells/well after 120 hours). This suggested that NAD(P)HX dehydratase, but not NAD(P)HX epimerase deficiency negatively impacts cellular health in the tested human cell line, leading to reduced cell proliferation and/or increased death rates, phenotypes that had not been observed in the yeast system.

**Figure 5:**
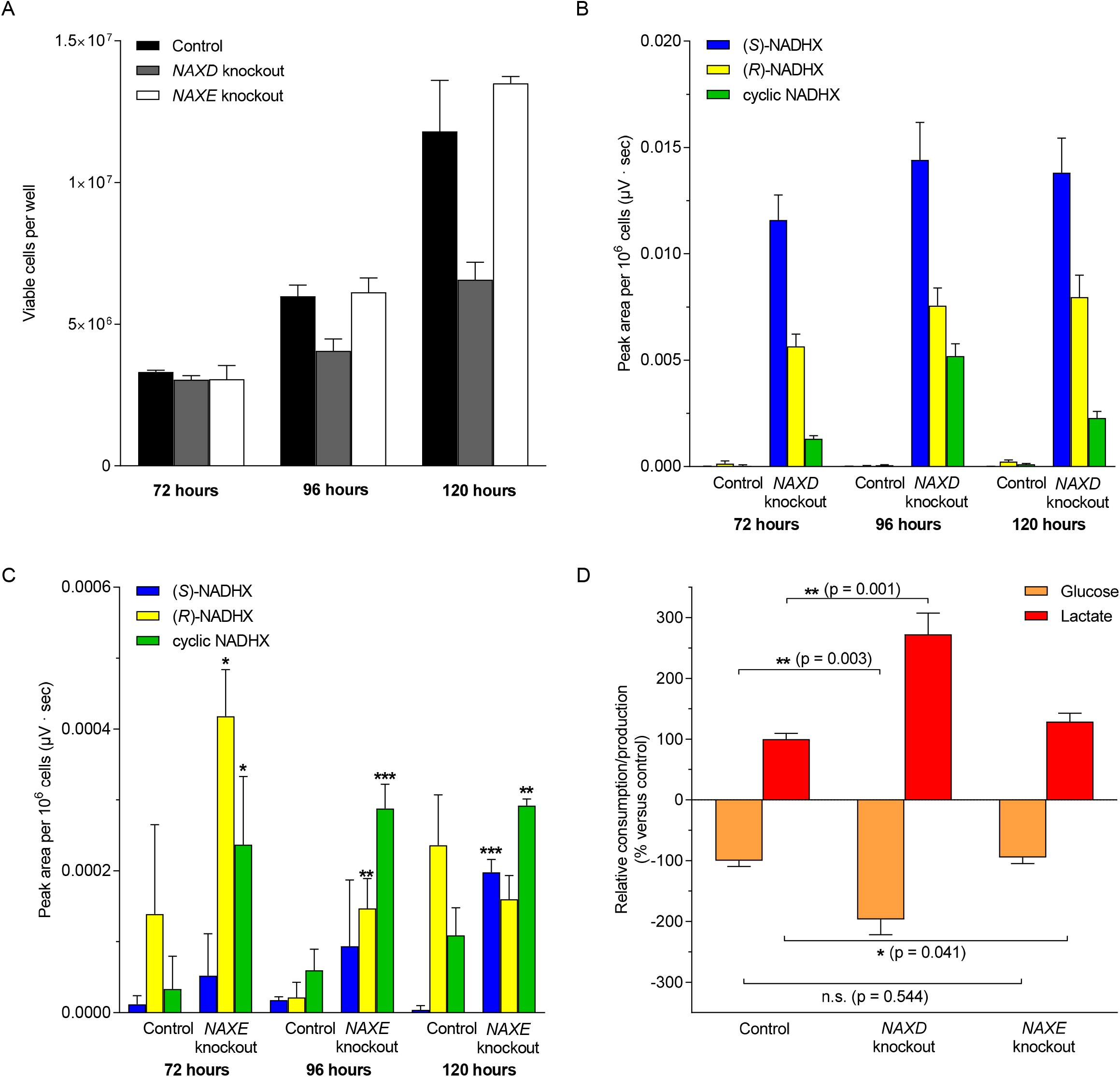
NADHX accumulation is accompanied by decreased cell viability, increased glucose uptake and increased lactate formation in a *NAXD* deficient human cell line. HAP1 control, *NAXD* (NAD(P)HX dehydratase) knockout, and *NAXE* (NAD(P)HX epimerase) knockout cell lines were cultivated for five days under standard conditions. (A) Cell number and cell viability were measured using a Nucleocounter NC-3000 at the indicated times after cell seeding. (B, C) Metabolites were extracted at the same time points to measure NADHX derivatives by HPLC-UV. Note the different scales used for the y-axis in panels (B) and (C) due to the much lower levels of NADHX detected in *NAXE* knockout cells. (D) Glucose consumption and lactate production, normalized to cell number, were calculated based on extracellular glucose and lactate concentrations and cell number increase measured between 72 and 96 hours after cell seeding. Relative values compared to control cell levels (glucose consumption and lactate production corresponded to 1.68 ± 0.11 mM/10^6^ cells and 2.60 ± 0.18 µmol/10^6^ cells, respectively) are shown. Consumptions and productions are represented as negative and positive values, respectively. All values shown are means and standard deviations of three biological replicates. Where indicated, statistically significant differences between knockout and control cell lines were determined applying an unpaired Student’s t-test and correcting for multiple comparisons using the Holm-Sidak method. All the NADHX concentrations shown in panel B for the *NAXD* knockout were significantly higher than the corresponding control cell line values (p ≤ 0.001; not indicated by asterisks).

Using our HPLC method, we also determined NADHX levels in metabolite extracts prepared 72, 96 and 120 hours after seeding the human cells. Analogous to the *ykl151c*Δ mutant, we found highly elevated levels of each NADHX derivative in the *NAXD* KO compared to the control cell line, but unlike for the yeast cells, the total amount of NADHX and the ratios of (*S*)-/(*R*)-NADHX versus cyclic NADHX seemed relatively independent from the sampling time point (Fig. 5B). (*S*)- and (*R*)-NADHX represented together more than 80% of the total NADHX measured at all time points and only minor amounts of cyclic NADHX were detected. A much more modest, but often significant accumulation of (*S*)-, (*R*)- and cyclic NADHX was also observed in the *NAXE* KO cell line compared to the control cell line (Fig. 5C). However, in contrast to the *NAXD* KO, cyclic NADHX alone accounted for more than 33% of the total NADHX and together with (*R*)-NADHX summed up to more than 70% of the total NADHX in the *NAXE* KO at least at the first two time points. Comparing the HPLC peak areas (normalized to the cell numbers), we estimated a more than 21-fold increase and a more than 3-fold increase for each of the NADHX derivatives in the *NAXD* and *NAXE* KO cell lines (except for (*R*)- NADHX after 120 hours in the *NAXE* KO, a measurement that was also not significant), respectively, compared to the control cells at each of the sampling points. At the 96 hour time point, this increase corresponded to more than 87-fold for the *NAXD* KO line. For the latter, using estimated intracellular volumes in our samples based on measurements of the average cell diameter at the respective sampling points with the NucleoCounter NC-3000, we calculated intracellular concentrations of about 51.0 (± 4.9) μM, 23.3 (± 2.4) μM, and 7.4 (± 0.7) μM (S)-, (*R*)-, and cyclic NADHX, respectively, after 72 hours of cultivation and of about 73.8 (± 4.3) μM, 37.4 (± 0.7) μM, and 30.0 (± 0.4) μM (S)-, (*R*)-, and cyclic NADHX, respectively, after 96 hours of cultivation (values are means ± SDs for n = 3 biological replicates). Due to the low peak areas detected for the control and *NAXE* KO cell lines, this type of absolute NADHX quantification could not be performed for these cell lines. The levels to which NADHX accumulated in our human NAD(P)HX dehydratase KO cells were in good agreement with the ones measured for the *ykl151c*Δ mutant. As opposed to the yeast system, however, a significant NADHX accumulation could also be detected in NAD(P)HX epimerase deficient human cells. Also, in contrast to our results in the *ykl151c*Δ mutant, we did not detect any significant changes in NAD^+^ levels in the NAD(P)HX repair deficient KO cell lines compared to the control cells (Fig. S4).

Additionally, in the cultivation conditions used for the human cell lines, we could so far not detect the decrease in intracellular serine levels that we had found in the yeast cells upon deletion of the NAD(P)HX dehydratase gene. It should be noted, however, that, while our yeast cells were cultivated in a minimal medium lacking serine, the medium used for the human cell cultivations had an initial serine concentration of about 360 μM. A potential inhibition of the endogenous 3PGA serine synthesis pathway by NADHX may therefore be compensated by increased uptake of extracellular serine in the human cell cultivation system. Alternatively, it could be that the human 3PGA dehydrogenase (PHGDH) is not inhibited by NADHX. Accordingly, we confirmed that human PHGDH acts as an NAD dependent dehydrogenase, as previously shown by others [43], but we also found that it lacks the transhydrogenase activity identified in the present study for the yeast homologs of the enzyme (Ser3 and Ser33). In preliminary spectrophotometric assays of the 3PGA dehydrogenase activity of human recombinant PHGDH (coupled to the PSAT1 reaction) based on monitoring NADH production at 340 nm, neither of the NADHX derivatives ((*S*)-, (*R*)-, or cyclic NADHX tested up to a concentration of 100 μM) inhibited this enzymatic activity when tested in the presence of 50 μM 3PGA and 50 μM NAD^+^ (data not shown).

Finally, as NADHX accumulation only became detectable in postdiauxic phase (i.e. upon depletion of extracellular glucose) in the yeast model, we determined the glucose and lactate concentrations in the medium of our human cell cultures. In control HAP1 cell cultivations, the extracellular glucose concentration dropped from an initial concentration of about 22.5 mM to about 12.6 mM (after 72 hours) and about 8.1 mM (after 96 hours). Strikingly, when normalizing glucose consumption between 72 and 96 hours to the cell number formed in this time interval, we observed a 2-fold higher glucose consumption for the *NAXD* KO cells than for the control and *NAXE* KO cells (Fig. 5D). In addition, we estimated a 2.7-fold higher lactate production for the *NAXD* KO cells than for the other two cell lines during that same period (Fig. 5D). These observations indicated that the relatively high intracellular NADHX levels accumulating in the absence of an NAD(P)HX dehydratase activity lead to an increased glycolytic flux, potentially to compensate for decreased ATP production due to mitochondrial dysfunction.

## DISCUSSION

### NAD(P)HX formation coincides with glucose depletion and slow proliferation rates in yeast

In this study, we show for the first time that NAD(P)HX can be formed in yeast cells and confirm that these damaged cofactors can also be formed in cultured human cells. These results, obtained in *S. cerevisiae* and the human near-haploid cell line HAP1, corroborate previous ones obtained in *A. thaliana* [10] and human fibroblasts [24]. While NADHX levels were low or barely detectable in wild-type cells, deletion of the NAD(P)HX dehydratase gene led to high accumulations of the NADH hydrates, reaching intracellular concentrations determined in our study to be in the 100-200 µM range in yeast and in human cells. Deletion of the NAD(P)HX epimerase gene led to more moderate, but detectable increases in NADHX levels in human cells, but not in our yeast model. It thus seems that, in the latter, the spontaneous epimerization rate between the (*S*)- and (*R*)-NADHX forms was not limiting efficient repair by the dehydratase alone. As spontaneous interconversion of the S- and R-epimers is highly accelerated by even a moderate decrease in pH from 7 to 6 *in vitro* [6], acidification during our yeast cultivations in non-buffered minimal media could be one reason for the dispensability of the epimerase under these conditions: the intracellular pH of yeast cells in comparable cultivations was found to be around 6.4 [44]. Using LC-MS, we also detected higher NADPHX levels in the yeast NAD(P)HX dehydratase deletion mutant *ykl151c*Δ. NADPHX detection remains, however, technically challenging (lower intracellular concentrations, lower stability, and earlier elution from HPLC column) and the remaining of our study focused on the NADHX derivative. As hydration damage affects the reduced forms of the nicotinamide dinucleotide cofactors and as metabolic regulations have evolved to maintain high NADPH/NADP ratios (while the opposite is true for NAD), it should be noted here that the analysis of NADPHX formation and its possible effects on cellular function remains a highly relevant topic that should be tackled in future studies.

Intriguingly, in the yeast mutants deficient in NAD(P)HX dehydratase, NADHX accumulation only became apparent in postdiauxic phase, a cultivation stage characterized by a switch from aerobic glucose fermentation to ethanol respiration and entry into quiescence. Also, while (*S*)- and (*R*)-NADHX were the most abundant derivatives in the NADHX dehydratase mutant *ykl151c*Δ in early postdiauxic phase, a further degradation product, cyclic NADHX, dominated in the *ykl151c*Δ metabolite profile in later stages. Several hypotheses may be formulated to explain the dynamics of NADHX formation observed here during liquid yeast cultivations. (1) Enzymes catalyzing the formation of NADHX as a side reaction may be upregulated in the postdiauxic phase. So far, GAPDH is the only enzyme for which such a side reaction has been described, based on *in vitro* assays [1], but it cannot be excluded that other NAD-dependent enzymes form NADHX as a side product as well. (2) In addition, spontaneous chemical conversion of NADH to NADHX may contribute more or less significantly to the increased damage load in that phase. It is well documented that NADHX formation is accelerated by low pH conditions [3, 7]. Thus, the medium acidification occurring during yeast flask cultivations [44], as already mentioned above in the context of NADHX epimerization, may progressively decrease the intracellular pH and thereby contribute to increased rates of spontaneous NADH hydration during postdiauxic phase. (3) Another factor that could contribute to the higher NADHX accumulation during postdiauxic phase is the transition from rapidly dividing cells to quiescent cells. Dilution by cell division may limit the intracellular accumulation of the damaged cofactor during exponential growth, despite the lack of the repair enzyme. As also suggested by others, damage dilution through cell division is probably a basic strategy to sustain cellular life despite the constant production of potentially harmful side products by all biological processes and may render rapidly dividing cells less dependent on damage control systems (such as metabolite repair enzymes) than non-dividing cells [45–47]. Expression data across several data sets, using the SPELL search engine [26], indicate higher expression of the *YKL151C* (and *YNL200C*) genes during diauxic shift and stationary phase, which further supports the notion that enzymatic repair of NAD(P)H damage may play a more critical role in quiescent cells and/or under respiratory conditions. In our standard experimental conditions, all tested yeast strains had doubling times of about 120 minutes, reflecting a rapid volume expansion of the cytoplasm. In contrast, the human HAP1 cells used presented doubling times of minimally eight hours, possibly explaining that in those cells NADHX detection was less dependent on the sampling time point. The damage dilution hypothesis could be tested using genetic systems in which the proliferative potential of daughter cells is eliminated and therefore leading to yeast cultivations enriched in aged mother cells [48, 49]. In such systems, NADHX may become detectable at earlier time points than in conventional cultivations. (4) Finally, as the metabolic state of yeast cells in exponential growth (glucose fermentation) is very different from the one in postdiauxic phase (ethanol respiration), it would also be interesting to study the dynamics of NADHX formation upon growth on respiratory carbon sources. Such experiments would be further motivated by recent findings showing an induction of *YKL151C* expression in yeast cells grown on glycerol as the sole carbon source [50]. Interestingly, a recent study on coenzyme longevity in microorganisms using a dynamic ^13^C labeling approach, completely independently concluded that repair of damaged cofactors (including NAD and NADP) may become ‘increasingly important the slower the cells divide’ [51].

### NAD(P)HX repair deficiency leads to discrete changes in gene expression and central carbon metabolism in yeast

Transcriptomic and metabolomic profiling of our NAD(P)HX repair deficient yeast strain allowed to uncover an association between NADHX accumulation and serine synthesis. The increase in intracellular NADHX concentrations during postdiauxic phase was correlated with a strong decrease in expression of a serine catabolic gene (*CHA1*) and a more than 50% decrease in intracellular serine levels. These observations motivated us to test the effect of NADHX on the enzyme catalyzing the first step of the main serine synthesis pathway in yeast, *i.e.* the only enzyme in this pathway reported to depend on the NAD cofactor. In our attempts to implement these tests, we made another discovery. Given that we were unable to detect the predicted phosphoglycerate dehydrogenase activity of the Ser3 and Ser33 enzymes and given some striking similarities between the *E. coli* pyridoxal phosphate synthesis pathway and the 3PGA-dependent serine synthesis pathway, we hypothesized that Ser3 and Ser33 may, as the related *E. coli* PdxB enzyme, catalyze a transhydrogenase reaction. We could indeed show, using LC-MS, that Ser3 and Ser33 oxidize 3PGA in the presence of α-ketoglutarate and in the absence of externally added NAD, with a concomitant production of 2-hydroxyglutarate, qualifying those enzymes as transhydrogenases rather than conventional dehydrogenases. In this study, we have not formally identified the configuration of the 2-hydroxyglutarate product. However, given our previous observation that Ser3 and Ser33 reduce αKG to the D enantiomer of this dicarboxylic acid [38], this is most likely also the product of the 3PGA-αKG transhydrogenase activity. While this work was ongoing, other investigators independently showed that the *Pseudomonas* and *E. coli* phosphoglycerate dehydrogenases (SerA) also actually act as 3PGA-αKG transhydrogenases that produce D-2HG [52, 53], perfectly corroborating our results with the yeast enzymes. Only relatively few examples of transhydrogenases are known so far. Interestingly, one of them is Dld3 (**EC 1.1.99.40**), a yeast enzyme for which we recently uncovered that it converts D-2HG to αKG while reducing pyruvate to D-lactate [38]. The Ser3/Ser33 and Dld3 transhydrogenase activities thus may intimately link serine biosynthesis to the respiratory chain via D-2HG metabolism.

Most relevant for the present study, we then went on to demonstrate that the transhydrogenase activity of Ser3 and Ser33 is potently inhibited by (*S*)- and (*R*)-NADHX and to a lesser extent by cyclic NADHX, a further degradation product of NADHX. These findings, obtained with the recombinant enzymes *in vitro*, strongly suggested that the serine depletion observed in the *ykl151c*Δ mutant resulted from inhibition of the main serine synthesis pathway at the level of the first dedicated step by the high intracellular NADHX concentrations. As our *in vitro* results would predict complete inhibition of 3PGA oxidation at the NADHX concentrations reached in the *ykl151c*Δ mutant in postdiauxic phase (more than 100-fold higher than the concentrations needed to completely block the Ser33 activity *in vitro*), the residual serine pool measured in the mutant most likely derived from the glyoxylate route for serine synthesis [37, 54]. Discovery of the extremely conserved, but apparently not essential, NAD(P)HX repair system [7], raised the question of its physiological relevance. While the recent description of patients presenting with a severe neurometabolic disorder caused by deficiency of this repair system unequivocally demonstrates its essential role in humans, our study provides a first answer in a more mechanistic direction to understand its biological role, pinpointing a metabolic blockage in a central metabolic pathway that would occur if NADHX was left to accumulate. It will be extremely interesting to further investigate the enzymatic properties of phosphoglycerate ‘dehydrogenases’ (transhydrogenase versus dehydrogenase activity; inhibition by small molecules) from different species across several domains of life to get a deeper understanding of how these properties relate to differences in protein sequence and structure. Ongoing experiments are designed to more deeply characterize the NAD(P)HX inhibition effects on yeast Ser3 and Ser33. Preliminary experiments indicate that the human phosphoglycerate dehydrogenase PHGDH does not catalyze a transhydrogenase reaction and that its NAD-dependent dehydrogenase activity is not inhibited by NADHX.

The Ser3/Ser33 reactions are located at a metabolic branch point, diverting the glycolytic intermediate 3PGA into serine synthesis, and therefore good targets for metabolic regulations. A possible feedback inhibition by serine at this step has already been reported [54]. Interestingly, co-regulated expression of *YKL151C* and its downstream neighboring gene *GPM1* (phosphoglycerate mutase converting 3PGA to 2PGA in glycolysis; **EC 5.4.2.11**) was found very recently [50]. This is mediated by a bidirectional intergenic promoter driving co-expression of *GPM1* as well as a *YKL151C* antisense transcript while repressing expression of the *YKL151C* sense mRNA, leading to anticorrelated *YKL151C* and *GPM1* transcript levels. During stationary phase and in the presence of non-fermentable carbon sources for instance, *GPM1* expression is repressed while *YKL151C* expression is induced [26, 50, 55]. One could imagine a protective role of Ykl151c for the serine synthesis pathway (by preventing NADHX accumulation), under conditions favoring low phosphoglycerate mutase activity and therefore potentially increased flux from 3PGA towards serine.

*CHA1*, a serine/threonine deaminase, was (after *YKL151C*) the gene with the most significantly changed expression (down-regulation) in the *ykl151c*Δ mutant compared to wild-type. This further supports that the serine synthesis pathway is impaired by NAD(P)HX accumulation. Expression of *CHA1* is indeed regulated by serine availability [41] and this has been shown to be mediated by sphingolipids [42]. *De novo* sphingolipid synthesis is initiated by reaction between serine and palmitoyl-CoA, followed by reduction to sphingoid bases. An increase in newly synthesized sphingoid bases triggers *CHA1* expression, leading to an increase in this serine catabolic enzyme, thereby forming a feedforward/feedback loop preventing sphingolipid accumulation by limiting serine levels. In the *ykl151c*Δ mutant, decreased *CHA1* expression may thus reflect lower sphingoid base levels due to decreased serine availability. More work will be needed to test for altered sphingolipid levels in the *ykl151c*Δ strain and potential associated phenotypes. Besides its function as a substrate for *de novo* sphingolipid synthesis, serine also serves as a proteinogenic amino acid, is a precursor for glycine, cysteine and phospholipids and feeds into one-carbon metabolism [56]. The latter is of particular interest given that one-carbon metabolism is essential for numerous cellular processes including protein, DNA, and RNA methylation as well as nucleotide biosynthesis [57]. In yeast, the mitochondrial and cytosolic serine hydroxymethyltransferases *SHM1* and *SHM2* catalyze the reversible conversion of serine to glycine with a concomitant conversion of tetrahydrofolate to 5,10-methylene-tetrahydrofolate, balancing the contribution of serine and glycine to one-carbon metabolism in both cellular compartments [58, 59]. Effects on other cellular processes due to the perturbation in serine synthesis in the *ykl151c*Δ mutant are therefore not unlikely and remain to be explored.

### NADHX accumulation affects cell viability in human cells, but not in yeast

Despite extensive growth profiling efforts, we could so far not identify conditions under which the *ykl151c*Δ mutant presents slowed growth. This is consistent with previous investigations, which also did not reveal any overt growth phenotypes for NAD(P)HX repair deficient mutants in *E. coli*, yeast, and *A. thaliana* [8, 10, 19, 20]. However, potassium limitation and ethanol/osmotic stress in an NAD(P)HX dehydratase knockout *Bacillus subtilis* strains led to prolonged lag phases and reduced viability [22], suggesting that this repair enzyme is beneficial for stress adaptation under certain conditions. Accordingly, the yeast NAD(P)HX dehydratase and/or epimerase are upregulated under several stress conditions such as heat stress, oxidative stress, and starvation [26, 50, 55]. In our human HAP1 cell line deficient in the NAD(P)HX dehydratase, we observed a decrease in cell viability upon prolonged cultivation already under standard conditions. While NAD(P)HX epimerase deficiency did not lead to decreased viability in our cell culture model, patients with mutations in the *NAXE* gene have recently been shown to suffer from a severe neurometabolic condition, triggered by febrile episodes and leading to early childhood death [23, 24].

In contrast to the yeast system, we measured NADHX accumulation in both the *NAXD* and *NAXE* HAP1 KO cells, but much higher levels were found in the *NAXD* KO cell line. In addition, *NAXD* KO cells, but not *NAXE* KO cells, showed higher glucose consumption and lactate formation. Elevated lactate levels and impaired mitochondrial function (decrease in complex I activity and pyruvate oxidation) were reported in the cerebrospinal fluid and in muscle biopsies, respectively, of *NAXE*-deficient subjects [24]. Fibroblast cell lines from these subjects also contained higher NADHX levels [24]. Taken together, our cell culture results and the results obtained with patient-derived material suggest that NAD(P)HX repair deficiency leads to perturbed mitochondrial energy metabolism (with a compensatory increase in glycolytic flux). While deficient NAD(P)HX epimerase activity has fatal consequences at the organismal level, the effects on cultured cell lines are more subtle ([24] and this study). In contrast, a loss of NAD(P)HX dehydratase activity leads to pronounced perturbations in cultured cells as described in this study. Mutations in this enzyme have not yet been reported in human subjects. The molecular mechanism linking NAD(P)HX repair deficiency and mitochondrial dysfunction remains to be elucidated; our NAD(P)HX KO cell line will be a useful model in that endeavor.

The onset of neurological symptoms in *NAXE*-deficient subjects upon fever episodes is consistent with the thermolability of NAD(P)H and the higher NAD(P)HX formation rates from NAD(P)H *in vitro* with increasing temperature [7]. Also in line with this, in our NAD(P)HX dehydratase KO yeast strain, cultivation at higher temperature led to increased NADHX accumulation and more pronounced NAD^+^ depletion. In the human NAD(P)HX repair deficient cells, we could not detect changes in NAD^+^ levels compared to control cells. It will be important to consolidate these results because a crucial question in view of finding treatment options for NAD(P)HX repair deficient subjects is whether the symptoms are mostly caused by a decrease of the functional pools of the NAD(P) cofactors or by toxic effects of the abnormal NAD(P)HX metabolites that accumulate. The former cause may be more straightforward to address, notably by supplementation of NAD precursor vitamins such as nicotinic acid, nicotinamide or nicotinamide riboside [60]. Unlike for the yeast model, we could not detect any changes in serine levels in our NAD(P)HX repair deficient human cell lines. However, as for the yeast model, combined transcriptomic and metabolomic profiling of the human *NAXD* KO cells may allow identification of metabolic blockages caused by NADHX or NADPHX and thereby guide further efforts to find treatment strategies for NAD(P)HX repair deficient subjects.

## MATERIALS and METHODS

### Materials

Reagents, of analytical grade whenever possible, were purchased from Sigma Aldrich (Bornem, Belgium), if not otherwise indicated.

### Generation of mutant and transformed yeast strains

The *Saccharomyces cerevisiae* strains used in this study are listed in Table S4. All strains were isogenic to the haploid reference strain BY4741, originating from the EUROSCARF collection (Frankfurt, Germany). The double knockout strain *ykl151c*Δ*ynl200c*Δ was generated by mating the corresponding haploid single mutant strains of opposing mating types, followed by sporulation on acetate-containing plates (10 g/l potassium acetate, 20 g/l agar) and tetrad dissection. Gene deletions in the knockout strains were confirmed by PCR analysis (see Table S5 for primer sequences).

The coding sequences of the *YKL151C*, *TDH1*, *TDH2*, and *TDH3* genes were PCR-amplified using primers containing attB sites (Table S5) and cloned into the pDONR221 vector (Invitrogen, Erembodegem, Belgium) using the BP clonase II enzyme mix (Invitrogen) according to the manufacturer’s instructions. The genes were then shuttled from the resulting Entry vectors to the pAG416-GPD-ccdB Destination vector (AddGene ID #14148) or the pAG426-GAL1-ccdB Destination vector (AddGene ID #14155) using the LR clonase II enzyme mix (Invitrogen) and the final Expression vectors were transformed into the desired yeast strains as described previously [38].

### Yeast cultivation

Yeast strains were cultivated in minimal defined medium (6.7 g/l Yeast Nitrogen Base (MP Biomedicals, Brussels, Belgium), 20 g/l D-glucose, 80 mg/l uracil, 80 mg/l L-methionine, 80 mg/l L-histidine, and 240 mg/l L-leucine), if not otherwise indicated. Batch cultures were incubated in flasks at 200 rpm and 30 °C, with a working volume corresponding to 15% of the flask volume or less. Liquid pre-cultures were started by inoculation from a single colony into rich YPD medium (10 g/l yeast extract, 20 g/l peptone, 20 g/l D-glucose) or SC medium (6.7 g/l YNB, 20 g/l D-glucose, 2 g/l SC amino acid mixture from MP Biomedicals) and incubated overnight at 30 °C. The next day, pre-cultures were diluted 1:50 in the medium used for the main culture and incubated for 6-8 hours. This second pre-culture served to inoculate the main culture at the desired optical density (OD). Cell growth was monitored by measuring OD600. Bio volume and cell concentration were determined using a Multisizer Z3 Analyzer (Beckman Coulter, Villepinte, France) after dilution in ISOTON^®^ II solution (Beckman Coulter).

### Human cell culture and viability assay

Control, *NAXD* knockout and *NAXE* knockout HAP1 cell lines were obtained from Horizon Discovery Group plc (Vienna, Austria) (Table S4). The provided knockout cell lines were generated by introducing a frameshift in the *NAXD* or *NAXE* genes using the CRISPR/Cas9 technology, resulting in an early truncation of the proteins. The genetic mutations were confirmed by Sanger sequencing (see Table S5 for primer sequences). Cells were maintained in IMDM medium supplemented with 10% fetal bovine serum and 1% penicillin-streptomycin (Thermo Fisher Scientific, Erembodegem, Belgium) at a constant CO2 level of 5% and at 37 °C. Cells were passaged after maximally 96 hours or before reaching a confluency of 85%. Medium was refreshed earlier if necessary. For cell culture experiments, cells were trypsinized (Thermo Fisher Scientific), counted (Nucleocounter NC-3000 from ChemoMetec, Kaiserslautern, Germany), and seeded (250,000 cells in 2 ml medium per well) into standard flat-bottom 6-well plates (Nunclon Delta Surface, Thermo Fisher Scientific). Culture medium was not changed until 96 hours after seeding, i.e. only for the last sampling point after 120 hours cells were supplied with new medium). Cell viability was determined using the Nucleocounter NC-3000 by mixing trypsinized cells at appropriate dilutions with a dye containing acridine orange and 4′,6-diamidin-2-phenylindol (Solution 13 AO – DAPI, ChemoMetec). The cell suspension was loaded onto NC-slides A8^TM^ (ChemoMetec) and measured with the “Viability and Cell Count Assay” adapted to the HAP1 cell type according to the manufacturer’s instructions.

### Yeast RNA extraction and gene expression analysis

Cells were grown in minimal defined medium until the desired growth phase. Culture aliquots (containing around 10^9^ cells for tiling arrays and around 2 × 10^8^ cells for qPCR experiments) were centrifuged for 5 min at 4,500 *g* and ambient temperature and the pellets immediately frozen at −80 °C. For RNA extraction, cells were resuspended in 200 μl of lysis buffer (50 mM TrisHCl, pH 7.0, 130 mM NaCl, 5 mM EDTA and 5% (w/v) sodium dodecyl sulfate) and transferred to a tube containing a 200 µl mixture of phenol-chloroform-isoamylalcohol (25:24:1) and about 500 µl of acid-washed glass beads. Lysis was performed in a Precellys cell homogenizer (VWR, Leuven, Belgium) for 20 sec at 5,000 rpm and ambient temperature. Following a 15-min centrifugation at 16,100 *g* and 4 °C, the upper aqueous phase containing the nucleic acids was transferred into a fresh reaction tube containing an equal amount of the phenol-chloroform-isoamylalcohol mixture. Samples were centrifuged and the previous step was repeated once more. Two further washing steps of the aqueous phase were performed using a chloroform-isoamylalcohol (24:1) mixture. RNA was precipitated from the washed aqueous phase by addition of a 3 M sodium acetate solution (1:20 of the total volume, Thermo Fisher Scientific) and two volumes of ethanol followed by an incubation at −20 °C for at least 30 minutes. After centrifugation for 15 min at 16,100 *g* and 4 °C, RNA pellets were washed with 80% ethanol, dried and resuspended in 50-150 µl RNase-free water. Samples were treated with ∼ 6 units DNase I per 25 µg RNA, using the TURBO DNA-free kit (Invitrogen) according to the manufacturer’s instructions. RNA was quantified using the Qubit RNA BR Assay Kit (Thermo Fisher Scientific).

For preparation of the tiling arrays, 20 µg RNA, 1.72 µg Random Primers (Invitrogen), and 0.034 µg Oligo(dT)12-18 Primers (Invitrogen) were incubated for 10 min at 70 °C and 10 min at 25 °C to reduce hairpin formation. First strand cDNA synthesis was performed in a mixture (60 µl) containing the denatured RNA and the primers, 1x reaction buffer from the SuperScript First-Strand Synthesis System (Invitrogen), 10 mM DTT, 0.5 mM dCTP/dATP/dGTP, 0.4 mM dTTP, 0.1 mM dUTP, 20 µg/ml Actinomycin D, and 30 U/µl SuperScript II Reverse Transcriptase (Invitrogen). The mixture was incubated for 60 min at 42 °C and for 10 min at 70 °C. RNA was degraded by addition of 3 µl of both RNaseH (30 Units, Epicentre, Madison, USA) and RNase Cocktail^TM^ (containing 30 Units RNase T1 and 1.5 Units RNase A, Thermo Fisher Scientific) and subsequent incubation at 37 °C and 70 °C for 20 min each. The remaining DNA was purified using the AMPure XP reagent (Beckman Coulter) according to the manufacturer’s instructions. For DNA fragmentation, biotinylation and labelling, the GeneChip WT Terminal labelling kit (Thermo Fisher Scientific) was used starting from 5.5 µg cDNA. Finally, labelled cDNA was hybridized to the Affymetrix *S. cerevisiae* tiling array (6.5 million 25-mer probes with an 8 nucleotide offset between each probe, PN 520055, Affymetrix, Thermo Fisher Scientific) using the GeneChip Hybridization, Wash and Stain kit (Affymetrix, Thermo Fisher Scientific). Scanning of the chips was performed with an Affymetrix 3000 7G scanner. For raw data treatment, transcription intensity signals were corrected for probe-specific parameters through normalization with genomic DNA from *Saccharomyces cerevisiae* S288c and probes were mapped to the yeast genome based on the annotation from Xu *et al.* [32, 33]. Tiling array probe intensities were aggregated to gene expression level as described in [61].

Two genes with highly significantly changed expression as indicated by the tiling arrays were validated by qRT-PCR. For this, cDNA was synthesized using 1.5 µg total RNA and random hexamer primers with the RevertAid H Minus First Strand cDNA Synthesis Kit (Invitrogen) according to the manufacturer’s instructions. The qRT-PCR reactions were performed in LightCycler 480 Multiwell plates (384 wells, Roche, Luxembourg, Luxembourg) using the iQ SYBR Green Supermix (BioRad, Temse, Belgium) with 3 pmol of each the gene-specific forward and reverse primers and 2 µl cDNA (1:20 dilution) in a total volume of 20 µl. The primers used are listed in Table S5. -Fold changes were calculated based on three biological replicates using the 2^−ΔΔ*CT*^ method and *ALG9* as the reference gene, which showed little variation between replicates as well as during time course experiments [62].

### Metabolite extraction and sample preparation for metabolite analyses

Yeast metabolite extracts were generated using two different methods. For targeted analysis of NAD derivatives only, a method adapted from Sporty *et al.* [25] was used. In our experiments, we adapted the volume of culture aliquots to be extracted to always correspond to a total of 5 OD units, assuming a linear correlation between biomass and OD. Cells were washed once with 2 ml of 0.9% (w/v) NaCl before resuspending them in 600 µl extraction buffer (50 mM ammonium acetate, pH 7.0). Cells were blasted for 2 × 30 sec with a 60 sec break in a Precellys Cell Homogenizer (VWR) connected to a Cryolys cooling unit (VWR) in 2-ml re-enforced bead-beating tubes (VWR) filled with glass beads to the meniscus of the cell suspension. After two additional washing steps of the glass beads and one of the cell debris with extraction buffer and acetonitrile (1:3 v:v) [25], extracts were transferred to tubes containing 6 ml chloroform and incubated for 20 min on ice to remove non-polar compounds. After centrifugation, the upper aqueous phase was freeze-dried in a LabConco lyophilizer (cold trap temperature −82 °C, chamber vacuum 0.002 mbar, VWR) and stored at −80 °C until analysis.

For a broader metabolite profiling (amino acids, organic acids, and NAD derivatives), samples were generated as described in [38] with two minor changes: the ratio between bio volume and extraction fluid was reduced to 1:50 (v:v) and the extraction buffer (50% methanol, 50% TE buffer at pH 7.0) and chloroform were bubbled with N2 to reduce spontaneous metabolite oxidation. The resulting filtered extracts were injected directly for LC-MS/MS analysis of amino acids and organic acids, but lyophilized and stored as described above for subsequent analysis of NAD derivatives by HPLC. In this approach, a Multisizer Z3 (Beckman Coulter, Villepinte, France) was used to determine cell concentration and single cell volume in the culture aliquots and this information was used to determine the total biovolumes and to calculate intracellular metabolite concentrations for each sample.

For metabolite extraction from adherent human cultured cells, the extraction method used in [38] was adapted as follows: cells were washed with 2 ml 0.9% (w/v) NaCl and then extracted by scraping them into 600 μl N2-bubbled extraction buffer (50% methanol, 50% TE buffer (10 mM TrisHCl, 1 mM EDTA, pH 7.0) while maintaining the 6-well plate on an ice-cold plate (CoolSink^®^ XT 96F, Biocision). Lysates were transferred to tubes containing 600 μl of ice-cold chloroform, followed by 30-min shaking at −20 °C and centrifugation. The upper polar phase was filtered into fresh tubes and stored at −80 °C until analysis.

### Metabolite analyses by HPLC and LC-MS/MS

To measure and quantify NAD^+^, NADH and NADHX derivatives ((*S*)-, (*R*)- and cyclic NADHX), a previously published HPLC-UV method was adapted [7]. Measurements were performed on a Shimadzu Nexera UHPLC system equipped with two LC-30AD pumps, a SIL-30AC cooled autosampler, and an SPD-M20A photo-diode array detector. Before analysis, dried samples were resuspended in 40-75 µl of 50 mM ammonium acetate, pH 7.0. Baseline separation of the compounds was achieved on a Polaris C18-A column (150 × 4.6 mm, 3.0 µm particle size, 180 Å pore size, Agilent, Diegem, Belgium) connected to a Metaguard Polaris C18-A pre-column (for 4.6 mm ID columns, Agilent) using 50 mM ammonium acetate, pH 7.0 as the mobile phase and acetonitrile (ACN) for compound elution according to the following gradient profile: 0 to 20 min, 0 to 5% ACN; 20 to 27 min, 5 to 90% ACN; 27-28 min, 90 to 0% ACN, and re-equilibration for 7 minutes at 0% ACN before injecting the next sample. Detection was performed at 279 nm for NAD^+^ and NADHX and 340 nm for NADH. External calibration curves with NAD^+^, NADH, and NADHX standards (0.7 to 100 μM) were used for quantification. The (*S*)-, (*R*)- and cyclic NADHX standards were prepared using a preparative HPLC method adapted from [7]. The NADH derivatives were separated on a Luna C18(2) column (250 × 10.0 mm, 5.0 µm particle size, 100 Å pore size, Phenomenex, Utrecht, Netherlands) at a constant flow rate of 4 ml/min with a mobile phase containing 10 mM ammonium acetate, pH 7.0 (elution gradient: 0 to 25 min, 0.1 to 5% ACN; 25 to 26 min, 5 to 0.1% ACN, and re-equilibration for 9 minutes at 0.1% ACN). Peak fractions containing the highest amounts of the NADHX derivative of interest were collected and dried in a lyophilizer after adjusting the pH to 8.0 with NaOH. Reconstituted fractions were used for the phosphoglycerate transhydrogenase inhibition assays in addition to serving as standards for the HPLC analyses. The (*S*)-, (*R*)- and cyclic NADHX concentrations in the standard solutions were determined spectrophotometrically using previously published extinction coefficients (ε_290_ = 13,500 M^−1^cm^−1^ for (*S*)- and (*R*)-NADHX and ε_290_ = 15,000 M^−1^cm^−1^ for cyclic NADHX [63]).

To quantify all 20 physiological amino acids, a targeted LC-MS/MS method was used. Measurements were performed on a Shimadzu Nexera XR LC 20AD coupled to a Sciex API4000 Qtrap mass spectrometer equipped with an electrospray ion source. Nitrogen was supplied by an NMG33 generator (CMC, Eschborn, Germany). Absolute concentrations were calculated based on external calibration curves prepared in 50% LC-MS-grade methanol. Measurement values were normalized to ^13^C-labelled internal standards extracted by the cold methanol method described above from yeast cells grown on ^13^C-glucose using isotope dilution mass spectrometry, except for glycine, alanine, cysteine, threonine, lysine, and aspartate (no ^13^C-standard available). Analyte separation was achieved by hydrophilic interaction chromatography using a SeQuant ZIC^®^-HILIC columm (150 × 2.1 mm, 3.5 µm particle size, 100 Å pore size, Merck, Overijse, Belgium) equipped with a SecurityGuard ULTRA pre-column (UHPLC HILIC for 2.1 ID Columns, Phenomenex) at a constant flow rate of 0.25 ml/min and a temperature of 25 °C. Target metabolites were eluted using the following gradient, where solvent A was water:acetonitrile:formic acid (98.9:1:0.1) and solvent B was acetonitrile:water:formic acid (98.9:1:0.1): 0 to 2 min, isocratic at 90% B; 2 to 7 min, 90 to 20% B; 7 to 10 min, 20 to 10% B; 10 to 12 min, isocratic at 10% B; 12 to 12.1 min, 10 to 90% B, followed by a re-equilibration for 8.9 minutes at starting condition. The autosampler was kept cold at 4 °C. 1 µl of sample was injected and mass spectral data was acquired in positive scheduled multiple reaction monitoring mode. Source settings and mass transitions are given in Table S6.

For analysis of metabolites connected to the serine synthesis pathway (3-phosphoserine, 3-phosphoglycerate, α-ketoglutarate, and 2-hydroxyglutarate), a second targeted LC-MS/MS method was applied using the same instrumental setup. Chromatographic separation was achieved on a C18EVO column (150 × 2.1 mm, 3.5 µm particle size, 100 Å pore size, Phenomenex) connected to a guard column (SecurityGuard ULTRA cartridge, UHPLC C18EVO for 2.1 mm ID columns, Phenomenex), keeping the column oven temperature at 55 °C and using a flow rate of 0.2 ml/min in gradient mode where solvent A was water:formic acid (99.9:0.1) and solvent B was methanol:formic acid (99.9:0.1), according to the following profile: 0 to 2 min, isocratic at 0% B; 2 to 4 min, 0 to 10% B; 4 to 6.5 min, 10 to 95% B; 6.5 to 10 min, isocratic at 95% B, 10 to 10.1 min, 95% to 0% B, followed by a re-equilibration of 9.9 minutes at starting condition. 2 µl of sample were injected and the autosampler was kept cold at 4 °C. Mass spectral data was acquired in negative multiple reaction monitoring mode. Source settings and mass transitions are given in the Table S6.

### Expression and purification of yeast Ser3 and Ser33 and of human phosphoserine transaminase (PSAT1) and 3-phosphoglycerate dehydrogenase (PHGDH)

The *SER3* and *SER33* coding sequences were cloned into bacterial expression plasmids as described previously [38]. The pET15b-PSAT1 and pET28-PHGDH expression vectors were kindly provided by Emile Van Schaftingen and Guido Bommer, respectively (de Duve Institute, Brussels). All plasmids were verified by Sanger sequencing and chemically transformed into *E. coli* BL21 (DE3) cells (Thermo Fisher Scientific). Bacterial cultivation and protein expression were carried out as previously described [38, 64]. The final isopropyl β-D-1-thiogalactopyranoside concentrations for induction of protein expression were 0.5 mM for PHGDH, 1 mM for PSAT1, and 0.1 mM for Ser3 and Ser33. Subsequent purification steps were performed on an ÄKTA protein purifier (GE Healthcare) using 1 ml HisTrap columns (GE Healthcare) for nickel-affinity purification and two HiTrap Desalting columns (5 ml, GE Healthcare) connected in series for desalting. Purified and desalted protein fractions were analyzed by SDS-PAGE and the protein concentration was measured using a Bradford assay (BioRad). Glycerol was added to the active fractions at a final concentration of 10% (v/v) before storage at −80 °C.

### Enzymatic activity assays

The transhydrogenase activity of Ser3, Ser33, and PHGDH was measured by incubating the enzyme in a reaction mixture containing 25 mM HEPES, pH 7.1, 1 mM MgCl_2_, 1 mM DTT, 5 μM pyridoxal-5’-phosphate, 50 μM 3-phosphoglycerate, 200-250 µM L-glutamate, 25 μM α-ketoglutarate, and 50-100 μg/ml PSAT at 30 °C, if not indicated otherwise. After a brief pre-incubation of the mixture at 30 °C, the reaction was started by adding the enzyme at a final concentration of 25-40 µg/ml for Ser33 and 20 μg/ml for PHGDH. After 5-10 minutes, aliquots were heat-inactivated at 95 °C for 5 minutes, filtered (4 mm diameter filters, regenerated cellulose, 0.2 µM pore size, Phenomenex) into glass vials and analyzed by LC-MS/MS as described above for the serine synthesis pathway intermediates. For inhibition assays, (*S*)-, (*R*)-, and cyclic NADHX as well as NAD^+^ were added to the reaction mixture to final concentrations between 0 and 25 μM. For these inhibition studies, a mock preparative HPLC without NADHX injection was run to collect fractions at the retention times of the different NADHX derivatives; those fractions were added instead of the corresponding NADHX standards to the reaction mixture for negative control assays. Enzymatic activities were calculated based on the time-dependent amount of 2-hydroxyglutarate formed; incubation times were chosen so as to remain within the linear product formation frame. Control reactions run in the absence of the 3PGA substrate were used for background correction of the activity measurements. All assays were performed in triplicate if not indicated otherwise.

3PGA dehydrogenase activity of human PHGDH was measured spectrophotometrically at a wavelength of 340 nm to follow NADH formation. The reaction mixture (total volume 200 µl) containing 25 mM Tris, pH 9.0, 1 mM DTT, 1 mM MgCl_2_, 5 µM pyridoxal-5’-phosphate, 1 mM L-glutamate, 100 mM KCl, 50 µM 3PGA, 50 µM NAD^+^, and 50 µg/ml PSAT was kept in a 96-well plate (Greiner Bio-One, Vilvoorde, Belgium) in a TECAN M200 Pro plate reader (Mechelen, Belgium) at 30 °C until a stable signal was reached and the reaction then started by the addition of 3 µg/ml PHGDH.

### Small molecule assays

Glucose and lactate levels in spent human cell culture media were measured using a YSI 2950D biochemistry analyzer (Yellow Springs Instruments, Yellow Springs, Ohio, USA) as described in [38] and according to the manufacturer’s instruction.

### Statistical analysis

An equal variance, unpaired Student’s t-test was applied to gene expression, metabolite concentration and LC-MS(/MS) signal intensity data to determine significant differences (n.s. = p ≥ 0.05; p < 0.05 = *, p < 0.01 = **, p < 0.001 = ***), if not indicated otherwise. For the differential pathway analysis of the tiling array results, a multiple hypothesis testing adjustment was additionally performed to increase the robustness of the statistical results.

## Acknowledgments

This work was supported by the Fonds National de la Recherche Luxembourg through an Inter-CORE grant (number 5773107) to CLL and LMS, an AFR PhD grant (number 4044610) to JBK and a CORE Junior grant (number 11339953) to NP as well as by the Fondation du Pélican de Mie et Pierre Hippert-Faber through a scholarship to JBK.

We thank Guido Bommer and Emile Van Schaftingen for providing the *ykl151c*Δ and *ynl200c*Δ mutant strains and expression plasmids for human PSAT1 and PHGDH. We thank Rita Gemayel for generating the *ykl151c*Δ*ynl200c*Δ double knockout strain. The pAG416-GPD-ccdB plasmid was a gift from Susan Lindquist’s laboratory (Addgene plasmid # 14148 and # 14155). We thank Andrew D. Hanson, Mona El-Badawi-Sidhu, and John Meissen for support with the LC-MS based analysis of NAD(P)HX derivatives in yeast extracts. We thank the EMBL Genomics Core Facility for technical support with the tiling array experiments. We thank Daniel Kay for technical help with cell culture viability assays and metabolome sampling from yeast and cell culture extracts. Finally, we thank Enrico Glaab for fruitful discussions on tiling array data analysis and for performing the differential pathway analysis.

## Author contributions

JBK and CLL designed the bulk of the experimental work and JBK implemented a majority of this work. JBK and CLL mainly analyzed and interpreted the data. JFC contributed to the metabolite extractions and the enzyme characterization experiments. CZ and LMS contributed the tiling arrays (on RNA samples prepared by JBK) and the corresponding raw data analysis. The gene expression data was further analysed by JBK and PPJ, who also contributed to growth profiling of mutant yeast strains. NP developed and validated the targeted LC-MS/MS methods and used them for amino acid and serine pathway intermediate analyses. OF contributed LC-MS-based confirmation of NAD(P)HX presence in yeast extracts. JBK and CLL wrote the manuscript. All authors revised and approved the final version of the manuscript.

## SUPPLEMENTAL MATERIAL

## SUPPLEMENTAL FIGURES

**Figure S1.**
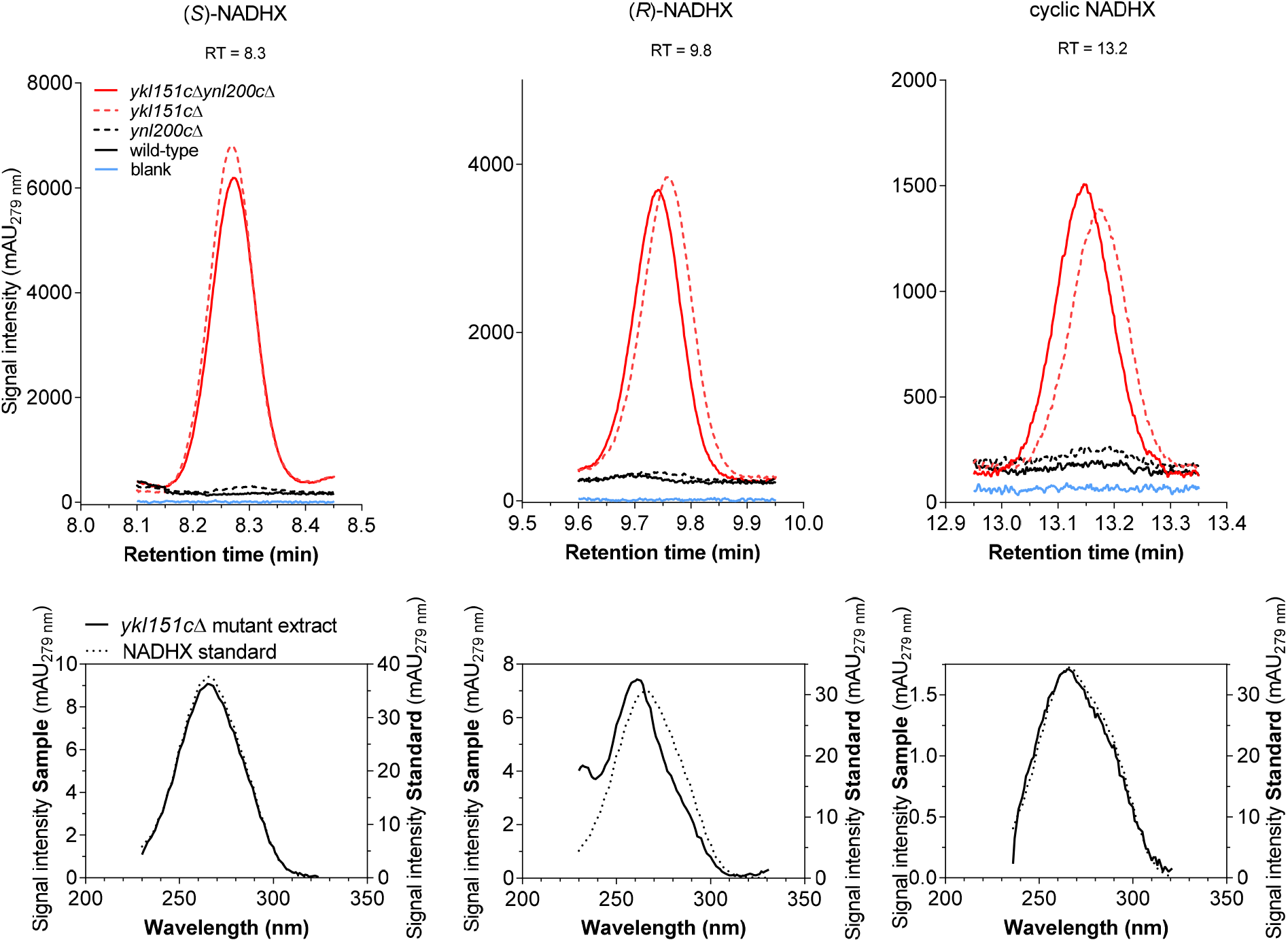
Identification of NADHX derivatives in yeast extracts by HPLC-UV. Wild-type (continuous black), NAD(P)HX epimerase deficient (*ynl200c*Δ, dashed black), and NAD(P)HX dehydratase deficient (*ykl151c*Δ, dashed red) yeast cells as well as cells deficient in both enzymes (*ykl151c*Δ*ynl200c*Δ, continuous red) were extracted around 1 hour into postdiauxic phase by the method adapted from Sporty *et al.* [1] as described in the Methods section of the main text. Cell extracts were analyzed by HPLC-UV and NADHX peaks were identified based on retention time and UV absorption spectrum in comparison to purified standards. For each strain, a representative chromatographic peak at the expected retention time is shown. A mobile phase ‘blank’ run (50 mM ammonium acetate, pH 7.0) is also shown as a background control (continuous blue line in upper panels). The UV absorption spectra of the NADHX derivatives in a *ykl151c*Δ mutant extract (dashed red line in upper panels) are overlaid with the spectra of the corresponding standards (lower panels). A contaminating compound present in yeast extracts in early postdiauxic phase co-elutes with (*R*)-NADHX, causing a slight shift of the absorption spectrum of the corresponding peak from the standard spectrum.

**Figure S2.**
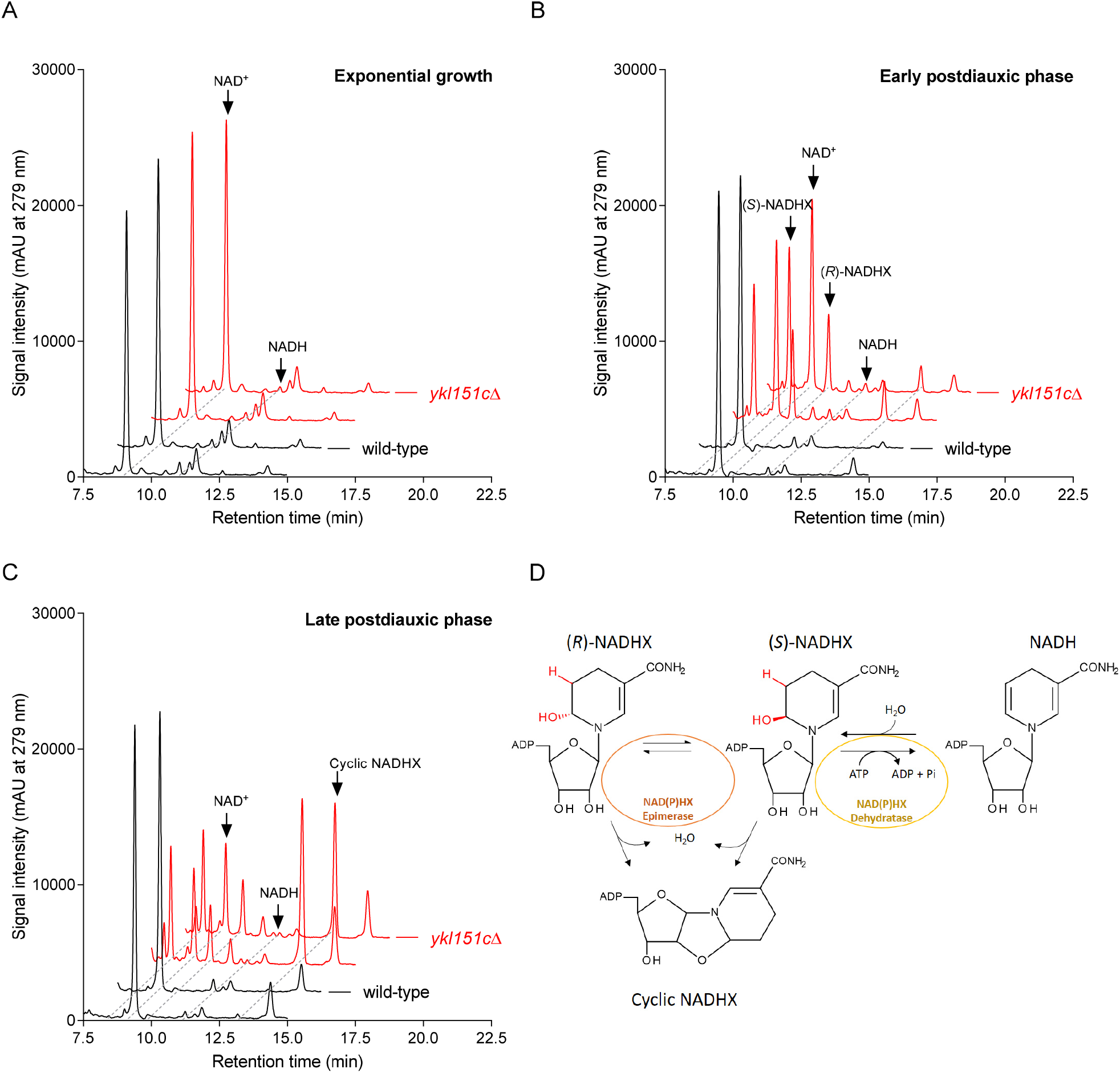
NADHX accumulation in yeast cells deficient in NAD(P)HX dehydratase. Wild-type (black) and NAD(P)HX dehydratase deficient (*ykl151c*Δ; red) yeast cells were extracted by the method adapted from Sporty *et al.* [1] during exponential growth (A), early postdiauxic phase (B), and late postdiauxic phase (C), as described in the Methods section of the main text. Cell extracts were analyzed by HPLC-UV and peaks were identified based on retention time and UV absorption spectrum comparison with appropriate standards. For each strain and growth phase, two representative HPLC chromatograms are shown. (D) The two NADHX epimers are formed by addition of a water molecule to the 5,6-double bond of the nicotinamide ring of NADH. If not repaired by NAD(P)HX dehydratase (yellow) and/or NAD(P)HX epimerase (orange), they undergo further degradation to cyclic NADHX.

**Figure S3.**
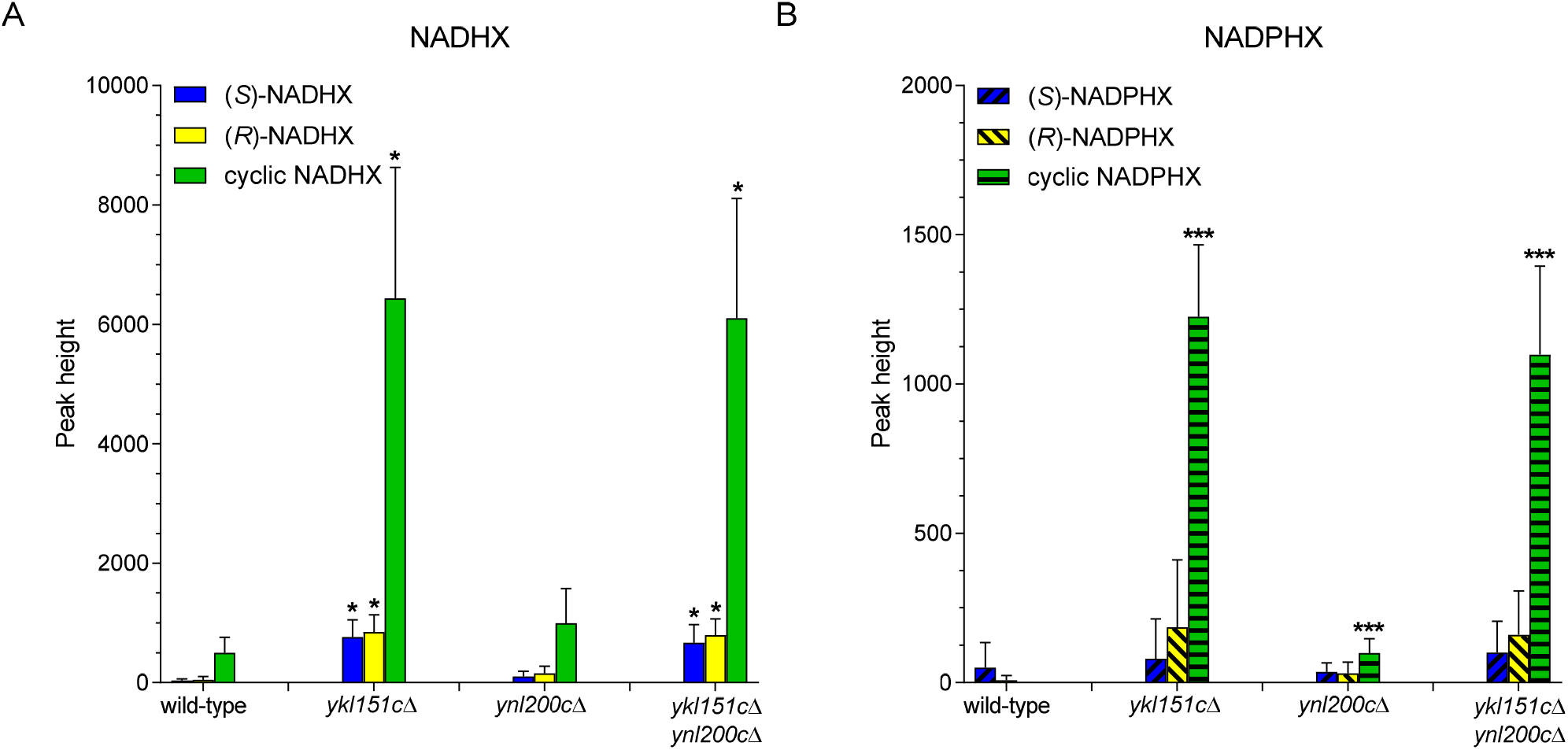
Identification of NADHX and NADPHX derivatives in yeast extracts by LC-MS. Wild-type, NAD(P)HX epimerase deficient (*ynl200c*Δ), and NAD(P)HX dehydratase deficient (*ykl151c*Δ), as well as yeast cells deficient in both enzymes (*ykl151c*Δ*ynl200c*Δ) were sampled around 1 hour into postdiauxic phase. Metabolites from ∼5 mg (wet weight) of cells were extracted and analyzed by LC-MS using retention time and accurate mass for compound identification as described in Niehaus *et al.* [2]. Graphs show means and standard deviations of 6 biological replicates. Statistical significance was determined using an unpaired Student’s t-test, correcting for multiple comparisons using the Holm-Sidak method. p-values were determined for all NADHX derivatives individually (compared to the wild-type) without assuming a consistent SD (*p < 0.05, **p < 0.01, ***p < 0.001).

**Figure S4.**
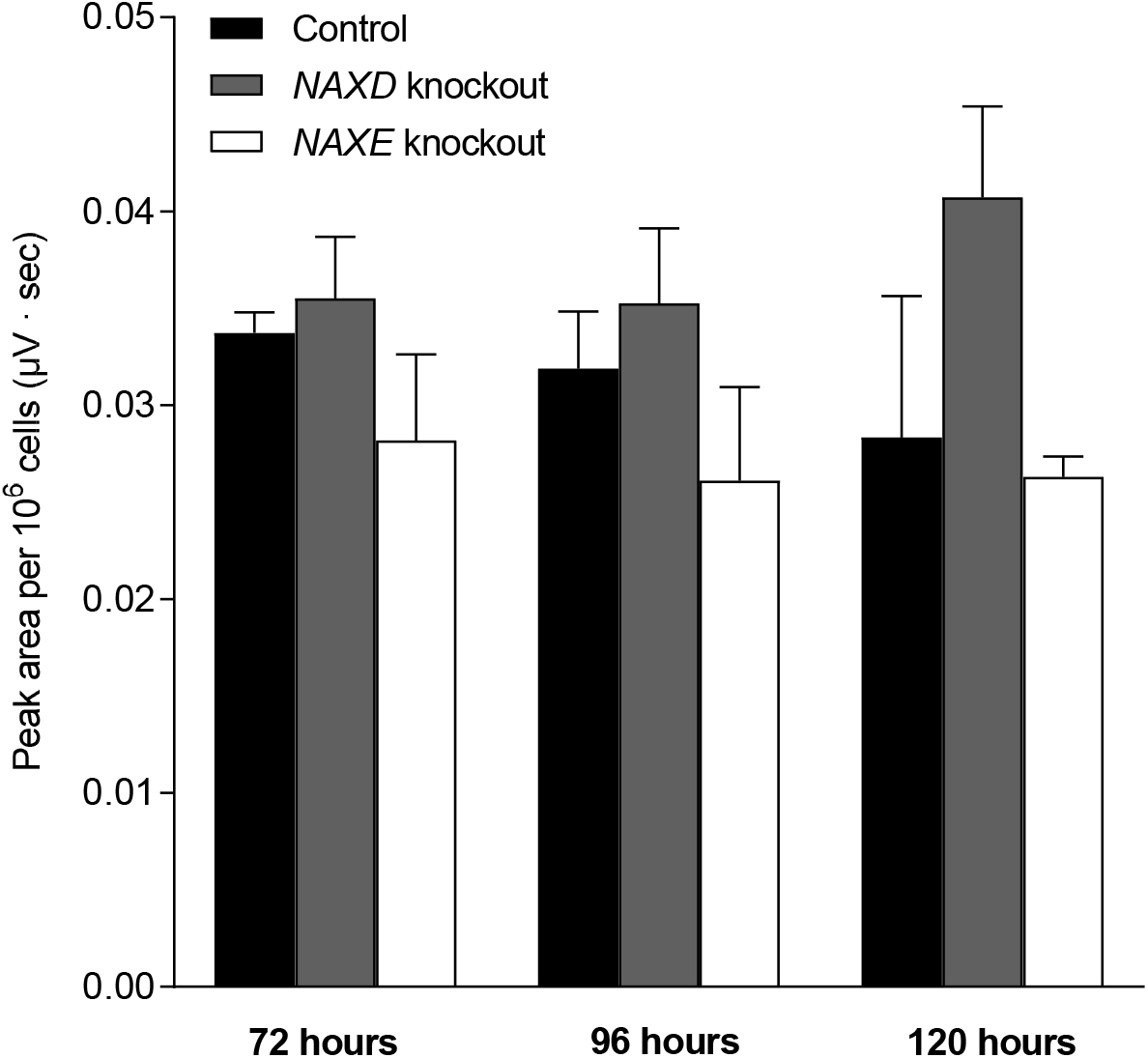
NAD(P)HX repair deficiency does not affect NAD^+^ levels in human HAP1 cells. HAP1 control, NAXD (NAD(P)HX dehydratase) knockout, and NAXE (NAD(P)HX epimerase) knockout cell lines were cultivated for five days under standard conditions. Metabolites were extracted at the indicated time points to measure NAD^+^ by HPLC-UV. All values shown are means and standard deviations of three biological replicates. Applying an unpaired Student’s t-test and correcting for multiple comparisons using the Holm-Sidak method, no significant differences between the control cell line and the NAD(P)HX repair deficient cell lines were found for NAD^+^ levels (p > 0.05).

## SUPPLEMENTAL TABLES

<u>Table S1</u> **Tested growth conditions for NAD(P)HX repair deficient yeast strains.** Wild-type and NAD(P)HX repair deficient strains (*ykl151c*Δ, *ynl200c*Δ, and/or *ykl151c*Δ*ynl200c*Δ) were cultivated under various conditions to screen for differences in biomass yield, growth rate, and/or duration of lag phase. Differences in fitness were evaluated using the following readouts: 1) Spotting assays, for which cultures were adjusted to the same OD and 4-5 serial dilutions were spotted on agar plates. Plates were maintained at 30 °C for several days until colonies became visible; 2) Growth curves obtained by plotting optical density measurements at 600 nm of liquid cultures obtained either manually in culture aliquots or semi-continuously in a TECAN M200Pro plate reader; 3) Growth parameters (length of lag phase, doubling time, and yield of biomass) obtained by pre-processing and analyzing OD600 measurements from 384 well-plate microcultivations with the GATHODE software as described in Jung *et al.* 2015 [3]. Standard minimal defined media contained 6.7 g/l YNB, 80 mg/l uracil, histidine, and methionine, 240 mg/l leucine, and 2% glucose, if not otherwise indicated. Rich media contained 2% peptone, 2% glucose, and 1% yeast extract, if not otherwise indicated. Solid media were prepared by adding up to 3% agar to liquid media. Culture media were supplemented with various chemicals or carbon sources other than glucose. Main cultures were inoculated with cells from exponential, postdiauxic or stationary phase and maintained at standard cultivation temperature (30 °C), except if otherwise indicated. When exposing a culture continuously to a non-standard temperature, the culture was inoculated from a pre-culture (30 °C) and kept during the whole cultivation at the indicated temperature. For heat shock experiments, the main culture (30 °C) was incubated up to 75 °C for up to 180 mins and then moved back to 30 °C for the remaining culture duration. Cultivation media were not buffered; where indicated, the starting pH was adjusted to a value below or above the standard one (pH ∼ 4.5). *SEE SEPARATE EXCEL SHEET*

<u>Table S2</u> **Genome-wide comparison of gene expression levels between an NAD(P)HX dehydratase knockout (*ykl151c*Δ) strain and the wild-type strain.** The wild-type and *ykl151c*Δ mutant strains were cultivated in minimal defined medium with 2% glucose and RNA was extracted in early and late stationary phase as described in the Methods section of the main text. RNA was reverse-transcribed and tilling arrays were performed according to the experimental pipeline also described within this section. Applying an equal variance, unpaired Student’s t-test, genes with significantly changed expression levels (p < 0.05) between the wild-type and mutant strains were identified. The expression levels for all the analyzed genes are shown in separate sheets for both sampling points (tp1 and tp2). CUT, Cryptic Unstable Transcript; KO, knockout; SUT, Stable Unannotated Transcript; tp, time point; WT, wild-type. *SEE SEPARATE EXCEL SHEET*

<u>Table S3</u> **Intracellular amino acid concentrations in an NAD(P)HX dehydratase knockout (ykl151cΔ) strain compared to the wild-type strain.** The wild-type and *ykl151c*Δ mutant strains were transformed with either an empty control plasmid or a rescue plasmid expressing the *YKL151C* gene and grown in minimal defined medium with 2% glucose, but lacking uracil. Metabolites were extracted at the indicated sampling points using the cold methanol extraction method and amino acids were analyzed using the targeted LC-MS/MS method described in the Methods section of the main text. Values shown are concentrations given in μM and represent means and standard deviations of three biological replicates. Statistical significance of the KO versus WT–fold changes were determined using an equal variance, unpaired Student’s t-test. *SEE SEPARATE EXCEL SHEET*

**Table S4.**
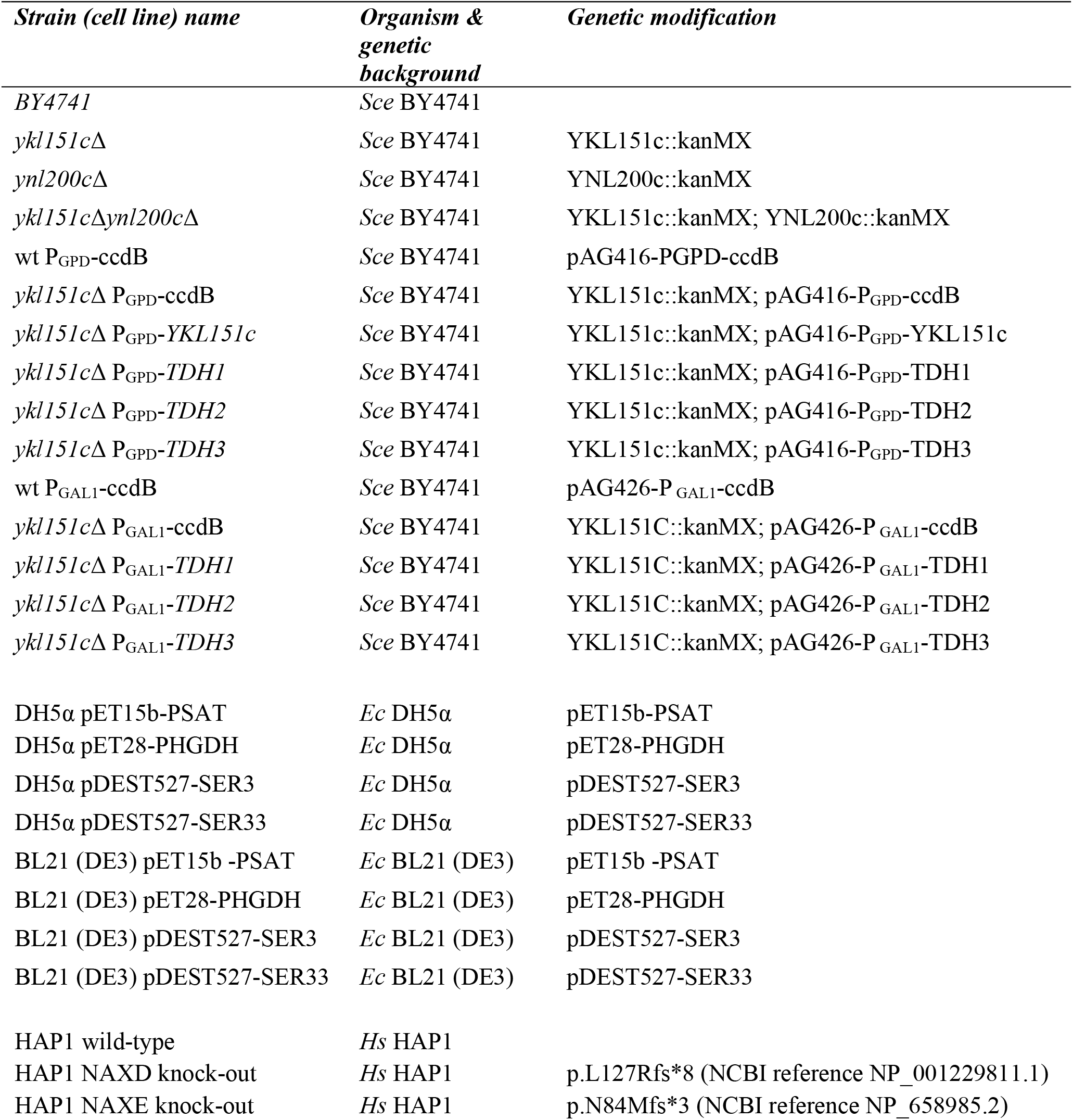
Yeast and bacterial strains and human cell lines used in this study. The yeast wild-type strain used for genetic modifications was the *Saccharomyces cerevisiae* BY4741 strain (*MAT***a**; *his3*Δ1; *leu2*Δ0; *met15*Δ0; *ura3*Δ0). The *Escherichia coli* BL21 (DE3) strain (F– ompT gal dcm lon hsdSB(rB− mB−) λ(DE3 [lacI lacUV5-T7 gene 1 ind1 sam7 nin5])) was used for protein expression and the *Escherichia coli* DH5α strain (*fhuA2 lac(del)U169 phoA glnV44 Φ80’ lacZ(del)M15 gyrA96 recA1 relA1 endA1 thi-1 hsdR17*) was used for plasmid propagation. HAP1 is a human near-haploid cell line derived from the male chronic myelogenous leukemia cell line KBM-7. ccdB, bacterial gene ensuring plasmid maintenance; *Ec, Escherichia coli*; *GAL1*, Galactokinase1; *GPD*, glyceraldehyde 3-phosphate dehydrogenase; *Hs, Homo sapiens;* kanMX, geneticin resistance cassette; *PHGDH,* phosphoglycerate dehydrogenase*; PSAT,* phosphoserine aminotransferase *Sce, Saccharomyces cerevisiae*; *TDH*, yeast glyceraldehyde-3-phosphate dehydrogenase.

**Table S5.**
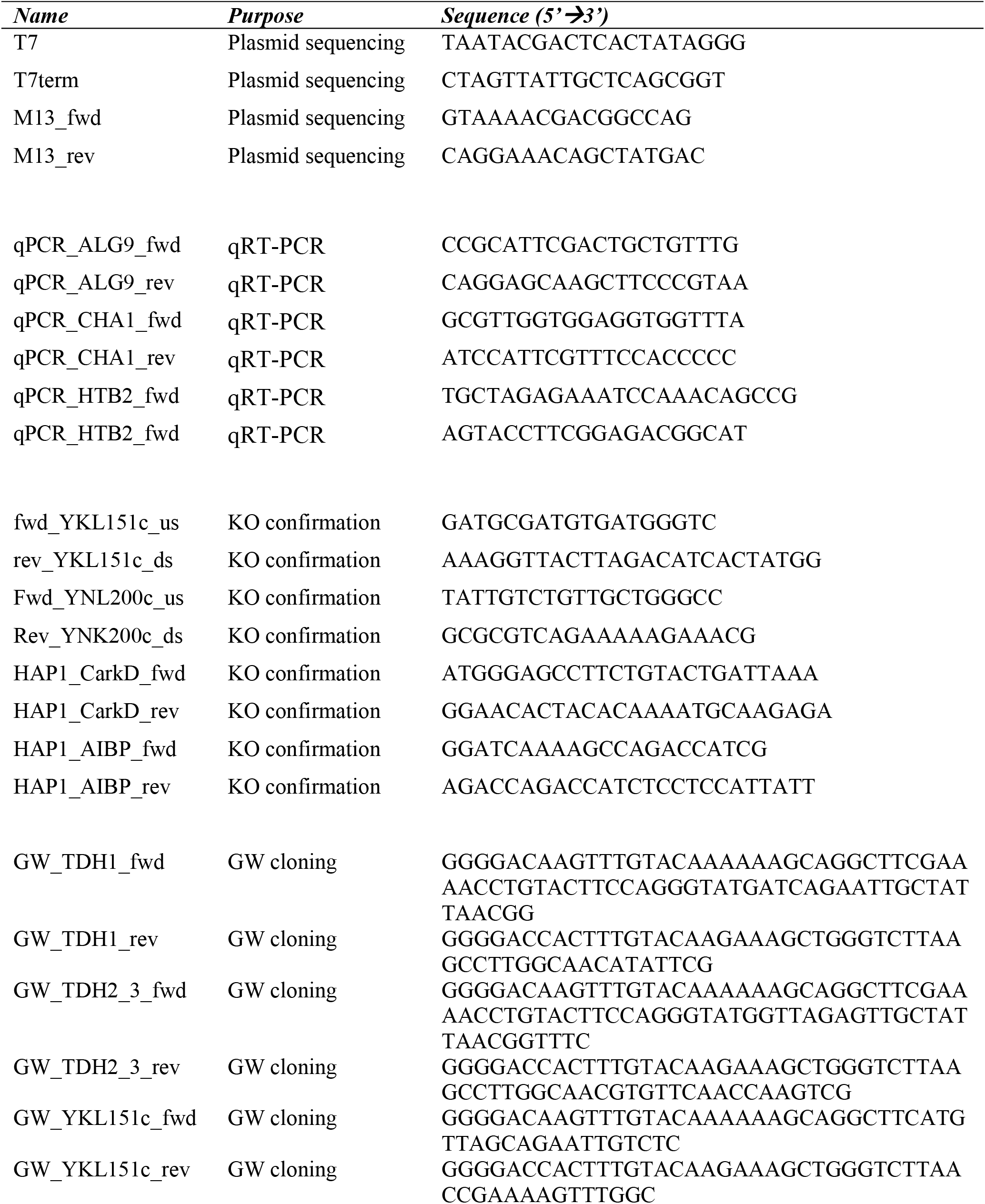
Primers used in this study for cloning, qRT-PCR and knockout confirmation. T7 and T7term were used to verify the bacterial expression plasmids pET15b–PSAT1 and pET28-PHGDH, and M13_fwd and M13_rev were used to verify the Entry Vectors, which served to create yeast expression strains (see Supplemental Table S4), by Sanger sequencing. fwd, forward; GW, Gateway; KO, knockout; rev, reverse.

**Table S6.** Source settings and mass transitions for the targeted LC-MS/MS methods used in this study. For the amino acid method, ionization parameters were set as follows: Nebulizer gas, 60 psi; auxiliary gas, 75 psi; curtain gas, 30 psi. Collision gas was set to medium. Ion spray voltage, 4500 V; temperature, 550 °C; interface heater, 100 °C. The MRM detection window was set to 30 sec. For the serine synthesis pathway intermediates method, ionization parameters were set as follows: Nebulizer gas, 60 psi; auxiliary gas, 75 psi; curtain gas, 30 psi. Collision gas was set to medium. Ion spray voltage, −4500 V; temperature, 550 °C; interface heater, 100 °C. The MRM detection window was set to 30 sec. Q1, mass detected in quadrupole 1 (mother ion); Q3, mass detected in quadrupole 3 (daughter ion); DP, declustering potential; EP, entrance potential; CE, collision energy; CXP, collision cell exit potential; RT, retention time; IS, internal standard.

| **Amino acid method** |  |  |  |  |  |  |  |
| --- | --- | --- | --- | --- | --- | --- | --- |
| **Metabolite** | **Q1**<br>**(m/z)** | **Q3**<br>**(m/z)** | **DP**<br>**(V)** | **EP (V)** | **CE (AU)** | **CXP (V)** | **RT (min)** |
| Glycine | 76.021 | 48 | 41 | 10 | 11 | 8 | 9.18 |
| Serine | 106.05 | 60.2 | 51 | 10 | 17 | 16 | 9.15 |
| Valine | 118.116 | 72.08 | 61 | 10 | 17 | 12 | 8.4 |
| Threonine | 120.088 | 74.2 | 56 | 10 | 17 | 12 | 8.99 |
| Leucine | 132.124 | 86.05 | 61 | 10 | 15 | 6 | 7.87 |
| Asparagine | 133.077 | 74.1 | 56 | 10 | 23 | 12 | 9.18 |
| Glutamine | 147.128 | 84 | 46 | 10 | 33 | 6 | 9.1 |
| Lysine | 147.168 | 84.2 | 56 | 10 | 23 | 14 | 10.45 |
| Histidine | 156.138 | 110 | 51 | 10 | 33 | 20 | 10.26 |
| Arginine | 175.091 | 70 | 56 | 10 | 35 | 12 | 10.47 |
| Tryptophan | 205.145 | 146 | 66 | 10 | 25 | 12 | 8.08 |
| Cysteine | 122.049 | 59.1 | 56 | 10 | 31 | 10 | 8.75 |
| Aspartate | 134.054 | 74 | 41 | 10 | 21 | 12 | 9.03 |
| Glutamate | 148.08 | 84 | 331 | 10 | 23 | 14 | 8.93 |
| Phenylalanine | 166.148 | 120.1 | 66 | 10 | 19 | 8 | 7.81 |
| Tyrosine | 182.139 | 136.2 | 56 | 10 | 19 | 8 | 8.52 |
| Alanine | 90.057 | 44.1 | 41 | 10 | 21 | 6 | 8.93 |
| Proline | 116.078 | 70 | 61 | 10 | 25 | 12 | 8.66 |
| Isoleucine | 132.096 | 86.2 | 51 | 10 | 15 | 6 | 7.99 |
| Methionine | 150.086 | 104.1 | 51 | 10 | 15 | 8 | 8.25 |
| Glycine IS | 78.04 | 49.1 | 41 | 10 | 11 | 8 | 9.18 |
| Serine IS | 109.11 | 62 | 51 | 10 | 17 | 16 | 9.15 |
| Valine IS | 123.06 | 76 | 61 | 10 | 17 | 12 | 8.4 |
| Leucine IS | 138.12 | 91.1 | 61 | 10 | 15 | 6 | 7.87 |
| Glutamine IS | 152.16 | 88.1 | 46 | 10 | 33 | 6 | 9.1 |
| Histidine IS | 162.12 | 115 | 51 | 10 | 33 | 20 | 10.26 |
| Arginine IS | 181.1 | 74.1 | 56 | 10 | 35 | 12 | 10.47 |
| Tryptophan IS | 216.07 | 198.9 | 66 | 10 | 25 | 12 | 8.08 |
| Aspartate IS | 138.07 | 76 | 41 | 10 | 21 | 12 | 9.03 |
| Glutamate IS | 153.14 | 88 | 331 | 10 | 23 | 14 | 8.93 |
| Phenylalanine IS | 175.1 | 128 | 66 | 10 | 19 | 8 | 7.81 |
| Tyrosine IS | 191.14 | 144.1 | 56 | 10 | 19 | 8 | 8.52 |
| Proline IS | 121.02 | 74.1 | 61 | 10 | 25 | 12 | 8.66 |
| Isoleucine IS | 138.12 | 91 | 51 | 10 | 15 | 6 | 7.99 |
| Methionine IS | 155.07 | 108 | 51 | 10 | 15 | 8 | 8.25 |

**Serine synthesis pathway intermediates method**
| **Metabolite** | **Q1**<br>**(m/z)** | **Q3**<br>**(m/z)** | **DP**<br>**(V)** | **EP**<br>**(V)** | **CE**<br>**(AU)** | **CXP**<br>**(V)** | **RT**<br>**(min)** |
| --- | --- | --- | --- | --- | --- | --- | --- |
| α-Ketoglutaric acid | 145 | 101 | -38 | -10 | -8 | -4 | 2.65 |
| 2-Hydroxyglutaric acid | 147 | 84.9 | -50 | -10 | -20 | -5 | 2.09 |
| 3-Phosphoglutaric acid | 185 | 96.7 | -40 | -10 | -20 | -5 | 2.22 |
| 3-Phosphoserine | 182.9 | 79.1 | -47 | -8 | -20 | -15 | 1.86 |

## Abbreviations

αKG: α-ketoglutarate
2HG: 2-hydroxyglutarate
GAPDH: glyceraldehyde-3-phosphate dehydrogenase
KO: knockout
*NAXE*: NAD(P)HX epimerase
*NAXD*: NAD(P)HX dehydratase
OD: optical density
3PGA: 3-phosphoglycerate
PHGDH: phosphoglycerate dehydrogenase
3PHP: 3-phosphohydroxypyruvate
3PSER: 3-phosphoserine
ORF: open reading frame
PSAT1: phosphoserine transaminase 1

